# An interaction between Gβγ and RNA polymerase II regulates transcription in cardiac fibroblasts

**DOI:** 10.1101/415935

**Authors:** Shahriar M. Khan, Ryan D. Martin, Sarah Gora, Celia Bouazza, Jace Jones-Tabah, Andy Zhang, Sarah MacKinnon, Phan Trieu, Paul B.S. Clarke, Jason C. Tanny, Terence E. Hébert

**Author notes:** To whom correspondence should be addressed. Dr. Terence E. Hébert, PhD, Department of Pharmacology and Therapeutics, McGill University, 3655 Promenade Sir-William-Osler, Room 1303 Montréal, Québec, H3G 1Y6, Canada OR Dr. Jason C. Tanny, PhD, Department of Pharmacology and Therapeutics, McGill University, 3655 Promenade Sir-William-Osler, Room 1303 Montréal, Québec, H3G 1Y6, Canada. These authors contributed equally to the study (Shahriar M. Khan, Ryan D. Martin).

## Abstract

Gβγ subunits are involved in many different signalling processes in various compartments of the cell, including the nucleus. To gain insight into the functions of nuclear Gβγ, we investigated the functional role of Gβγ signalling in regulation of GPCR-mediated gene expression in primary rat neonatal cardiac fibroblasts. Following activation of the angiotensin II type I receptor in these cells, Gβγ dimers interact with RNA polymerase II (RNAPII). Our findings suggest that Gβ_1_ recruitment to RNAPII negatively regulates the fibrotic transcriptional response, which can be overcome by strong fibrotic stimuli. The interaction between Gβγ subunits and RNAPII expands the role for Gβγ signalling in cardiac fibrosis. The Gβγ-RNAPII interaction was regulated by signaling pathways in HEK 293 cells that diverged from those operating in cardiac fibroblasts. Thus, the interaction may be a conserved feature of transcriptional regulation although such regulation may be cell-specific.

## INTRODUCTION

In recent years, study of the role of paracrine interactions between cardiomyocytes and cardiac fibroblasts in modulating the response to cardiac damage has expanded dramatically. Cardiac fibroblasts, in particular, respond dynamically following damage to the myocardium which is characterized by differentiation into myofibroblasts, increased proliferation and migration to areas of damage (Travers et al., 2016, Fu et al., 2018, Dobaczewski et al., 2010). This fibrotic response is modulated by the renin-angiotensin system, acting predominantly through the peptide ligand angiotensin II (Ang II) (Murphy et al., 2015, Kawano et al., 2000). Ang II drives changes in fibroblast function both directly and indirectly by increasing expression of other pro-fibrotic growth factors, such as transforming growth factor β1 (TGF-β1) (Campbell and Katwa, 1997). Collectively, these factors regulate alterations in cardiac architecture required for tissue repair by modulating the expression of genes encoding extracellular matrix proteins and proteases (Rosenkranz, 2004, Gao et al., 2009). Ang II also promotes cytokine secretion, thereby triggering autocrine and paracrine signalling to elicit further responses (Cheng et al., 2003, Ahmed et al., 2004). These signalling events create a feedforward loop, amplifying the fibrotic response from the initial area of damage to more distal regions of the heart (Ma et al., 2018). While the process initially aids in wound healing, a prolonged, activated fibrotic response worsens adverse cardiac remodelling and accelerates progression to heart failure (Travers et al., 2016, Weber et al., 2013). Importantly, inhibiting aspects of the fibrotic response reduces adverse cardiac remodelling (Fu et al., 2018, Weber and Diez, 2016). Hence, deciphering how Ang II signalling regulates pro-fibrotic gene expression is an important step towards understanding how these processes might be targeted therapeutically.

Cardiac fibroblasts respond to increased Ang II levels through Ang II type I (AT1R) and type II (AT2R) G protein-coupled receptors (GPCRs). Of these, the AT1R is responsible for positively regulating the fibrotic response in cardiac fibroblasts (Travers et al., 2016). The AT1R couples to multiple heterotrimeric G proteins composed of specific combinations of Gα and Gβγ subunits (Namkung et al., 2018). G proteins serve as signal transducers to relay extracellular ligands bound to GPCRs into activation of different intracellular signalling pathways (Khan et al., 2013). Gβγ subunits, like the more extensively studied Gα subunits, modulate a wide variety of canonical GPCR effectors at the cellular surface such as adenylyl cyclases, phospholipases and inwardly rectifying potassium channels (Khan et al., 2013, Dupré DJ, 2009, Smrcka, 2008). However, compared with Gα-mediated events, Gβγ-mediated signalling is relatively understudied and is complicated by the existence of 5 Gβ and 12 Gγ subunits which can combine in multiple ways to form obligate dimers. Gβγ subunits also regulate a variety of non-canonical effectors in distinct intracellular locations, and a number of studies have described roles for Gβγ signalling in the nucleus (Khan et al., 2013, Campden et al., 2015a). Nuclear Gβγ subunits modulate gene expression through interactions with a variety of transcription factors, such as adipocyte enhancer binding protein 1 (AEBP1), the AP-1 subunit c-Fos, HDAC5 and MEF2A (Park et al., 1999, Robitaille et al., 2010, Spiegelberg and Hamm, 2005, Bhatnagar et al., 2013). Furthermore, we have detected Gβ_1_ occupancy at numerous gene promoters in HEK 293 cells (Khan et al., 2015). While canonical Gβγ signalling has been implicated in both cardiac fibrosis and heart failure (Kamal et al., 2017, Travers et al., 2017), how nuclear Gβγ signalling impacts these events is currently unknown.

Here, we describe a novel interaction between Gβγ subunits and RNA polymerase II (RNAPII) which regulates the cardiac fibrotic response to Ang II activation of AT1R. We characterize the GPCR-dependent, signalling pathway-specific regulation of this interaction in primary neonatal rat cardiac fibroblasts and in HEK 293 cells. To understand the potential role of individual Gβγ subunits, we knocked down Gβ_1_ and Gβ_2_ as exemplars of Gβ subunits highly expressed in these cells and characterized how nuclear Gβ_1_, in particular, is a key regulator of AT1R-driven transcriptional changes.

## RESULTS

### Gβγ interaction with RNAPII following activation of Gαq-coupled GPCRs

As Gβγ interacts with transcription factors and occupies gene promoter regions, we hypothesized that Gβγ subunits interact with a protein complex ubiquitously involved in transcription, and we initially focused on RNAPII. We assessed the potential Gβγ-RNAPII interaction following endogenous M3-muscarinic acetylcholine receptors (M3-mAChRs) activation with carbachol in HEK 293F cells. An initial co-immunoprecipitation time course experiment revealed a carbachol-induced interaction between endogenous Gβγ subunits (Gβ_1-4_ detected with a pan-Gβ antibody) and Rpb1, the largest subunit of RNAPII, peaking between 45 and 120 mins (Supplemental Figure 1A, B). Immunoprecipitation of Rpb1 with two different antibodies also co-immunoprecipitated Gβ_1-4_ in an agonist-dependent manner (Supplemental Figure 1C). Further, we observed no basal or carbachol-dependent interaction of Rpb1 with Gαq/11 or ERK1/2 (Supplemental Figure 1D, E) suggesting that Gβγ was not in complex with these proteins when it was associated with RNAPII in the nucleus. Under similar conditions, we observed no basal or carbachol-dependent interaction of Gβγ subunits with the A194 subunit of RNA polymerase I (Supplemental Figure 1F), suggesting Gβγ is not recruited to all RNA polymerases.

We next assessed the whether the Gβγ-RNAPII interaction also occurred in primary rat neonatal cardiac fibroblasts following treatment with Ang II. A time-course co-immunoprecipitation experiment revealed an agonist induced Gβγ-RNAPII interaction with a major peak interaction observed 75 minutes post stimulation (**Figure 1A, B**). As cardiac fibroblasts express both AT1R and AT2R, we next examined which receptor subtype regulated the response, by pre-treatment with the AT1R-specific antagonist losartan. Pre-treatment of cells with losartan prior to Ang II treatment abolished the agonist-induced interaction, but preserved the basal interaction, suggesting that AT1R, and not AT2R, is primarily responsible for mediating the interaction (**Figure 1C, D**).

**Figure 1.**
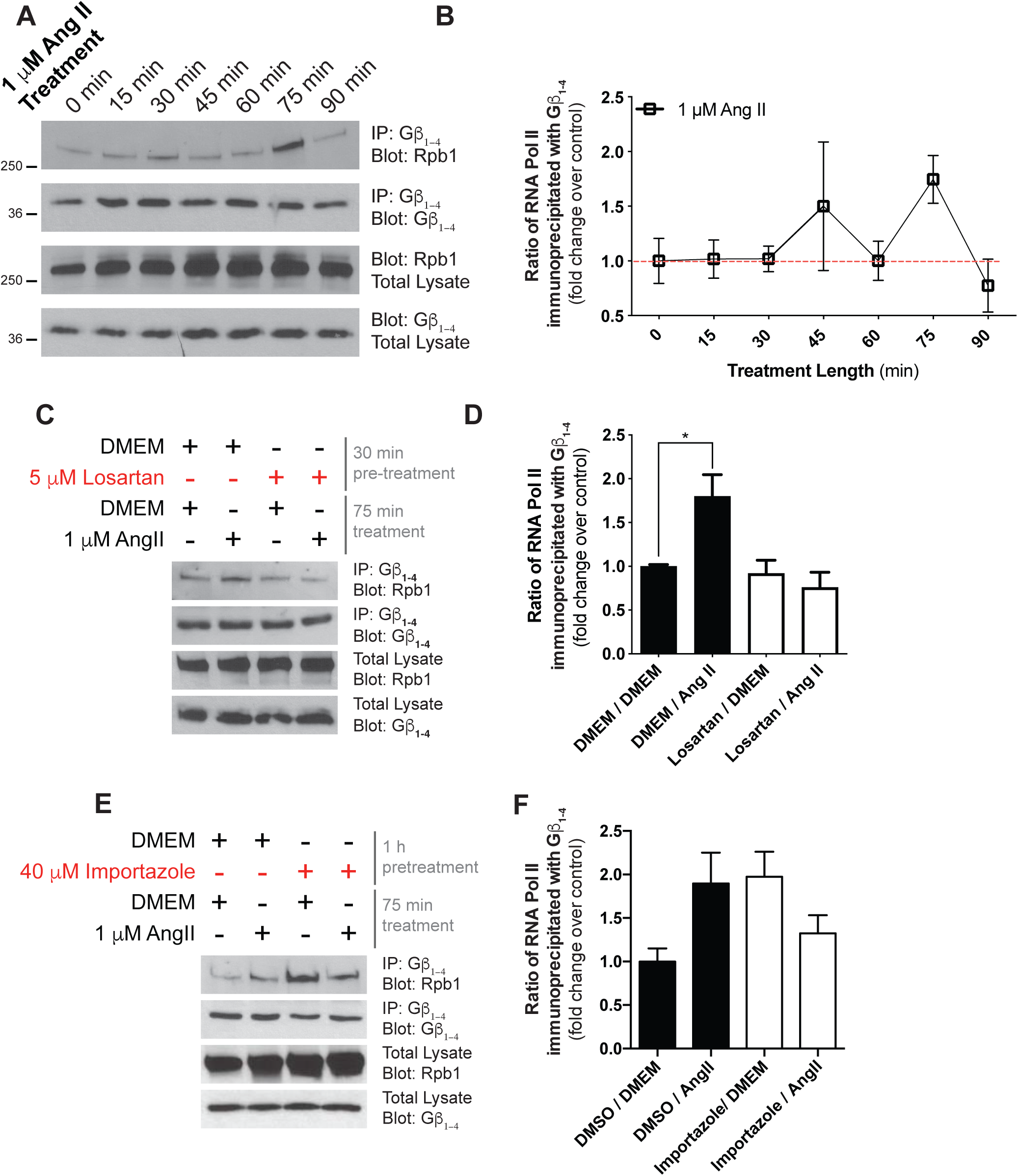
Characterization of Gβγ-RNAPII in rat neonatal cardiac fibroblasts. **(A)** Time course of the Ang II-stimulated interaction between Gβγ and Rpb1. The ratio of Rpb1 co-immunoprecipitated with Gβ_1-4_ upon treatment of 1 µM Ang II treatment at the indicated timepoints in cardiac fibroblasts was assessed. **(B)** Densitometry-based quantification of panel A was used to determine the ratio of Rpb1 to Gβ_1-4_ immunoprecipitated at each time point. The fold change over the 0 min time point was then calculated. Data is representative of four independent experiments. **(C)** Effect of AT1R antagonist losartan pre-treatment on the Ang II-mediated interaction, demonstrating angiotensin receptor subtype selectivity. **(D)** Densitometry-based quantification of AT1R antagonist effect on Ang II-induced interaction. The ratio of Rpb1 immunoprecipitated with Gβ_1-4_ was determined for each condition and fold change over DMSO/DMEM was calculated. Data is representative of three independent experiments. **(E)** Assessment of the necessity of Gβγ import into the nucleus for interaction to occur upon AT1R stimulation with Ang II. Cardiac fibroblasts were pretreated for 1 h with importazole prior to Ang II stimulation. Data are representative of four independent experiments. **(F)** Densitometry-based quantification of the Ang II induced interaction and the effect of nuclear import inhibition. The ratio of Rpb1 to Gβ_1-4_ immunoprecipitated was determined and normalized to fold change over DMSO/DMEM treatment. In all panels, data represents mean ± S.E.M, * indicates p<0.05, ** indicates p<0.01.

Although several Gβγ isoforms have been detected in the nucleus (Bhatnagar et al., 2013, Campden et al., 2015b, Zhang et al., 2001), the mechanisms leading to entry of Gβγ into the nucleus remain unknown. Using subcellular fractionation following M3-mAChR activation in HEK 293F cells, we observed importin-β dependent translocation of Gβγ into the nucleus **(data not shown).** In addition, the agonist-dependent interaction of Gβ_1-4_ and RNAPII was blocked by importazole pre-treatment, suggesting that nuclear import of Gβ_1-4_ is required for the interaction with RNAPII in these cells (Supplemental Figure 2A, B). Next, we determined the effect of importazole pre-treatment on the Ang II-mediated Gβγ-RNAPII interaction in cardiac fibroblasts. The Gβγ-RNAPII interaction was also ablated when nuclear import via importin-β was inhibited, suggesting again that Gβγ subunits must translocate to the nucleus for the interaction with RNAPII to occur (**Figure 1E, F**).

### Signalling pathways regulating Gβγ-RNAPII interaction are cell-specific

We next examined signalling events downstream of receptor activation that could mediate the interaction between Gβγ subunits and RNAPII. To this end, we pursued a pharmacological and genetic approach using both cardiac fibroblasts (**Figure 2**) and HEK 293F cells (**Figure 3**). Our data indicated that the pathways responsible for promoting the Gβγ-RNAPII interaction are cell type specific. Since AT1R couples to both Gq/11 and Gi/o G proteins (Sauliere et al., 2012), we used FR900359 to inhibit Gαq/11 (Schrage et al., 2015) and pertussis toxin (PTX) to inhibit Gαi/o. The agonist-induced response was markedly (∼80%) decreased by the Gαq/11 inhibitor, and also decreased (∼30%) by the Gαi/o inhibitor, demonstrating that AT1R signalling through Gαq is the primary pathway leading to increased Gβγ-RNAPII interaction (**Figure 2A-B**, Supplemental Figure 3A-B). We next used U71322 to inhibit the activity of phospholipase Cβ (PLCβ), downstream of both Gq/11 and Gi/o (the latter via Gβγ signalling). In cardiac fibroblasts, pre-treatment of U71322 blocked the agonist-induced Gβγ-RNAPII interaction with no effect on the basal interaction, suggesting a pivotal role for PLCβ (**Figure 2C**, Supplemental Figure 3C). Chelation of Ca^2+^ using BAPTA-AM in cardiac fibroblasts also abrogated the Ang II-induced Gβγ-RNAPII interaction (**Figure 2D**, Supplemental Figure 3D), as did treatment with the PKC inhibitor Gö6983 and the CaMKII inhibitor KN-93 (**Figure 2E, F**, Supplemental Figure 3E, F). Conversely, the MEK1 inhibitor U0126 led to an increased basal Gβγ-RNAPII interaction but abrogated the Ang II-induced interaction (**Figure 2G**, Supplemental Figure 3G). Lastly, the calcineurin inhibitor cyclosporin A lead to an increased basal interaction did not prevent further Ang II-dependent increase in interaction (**Figure 2H**, Supplemental Figure 3H).

**Figure 2.**
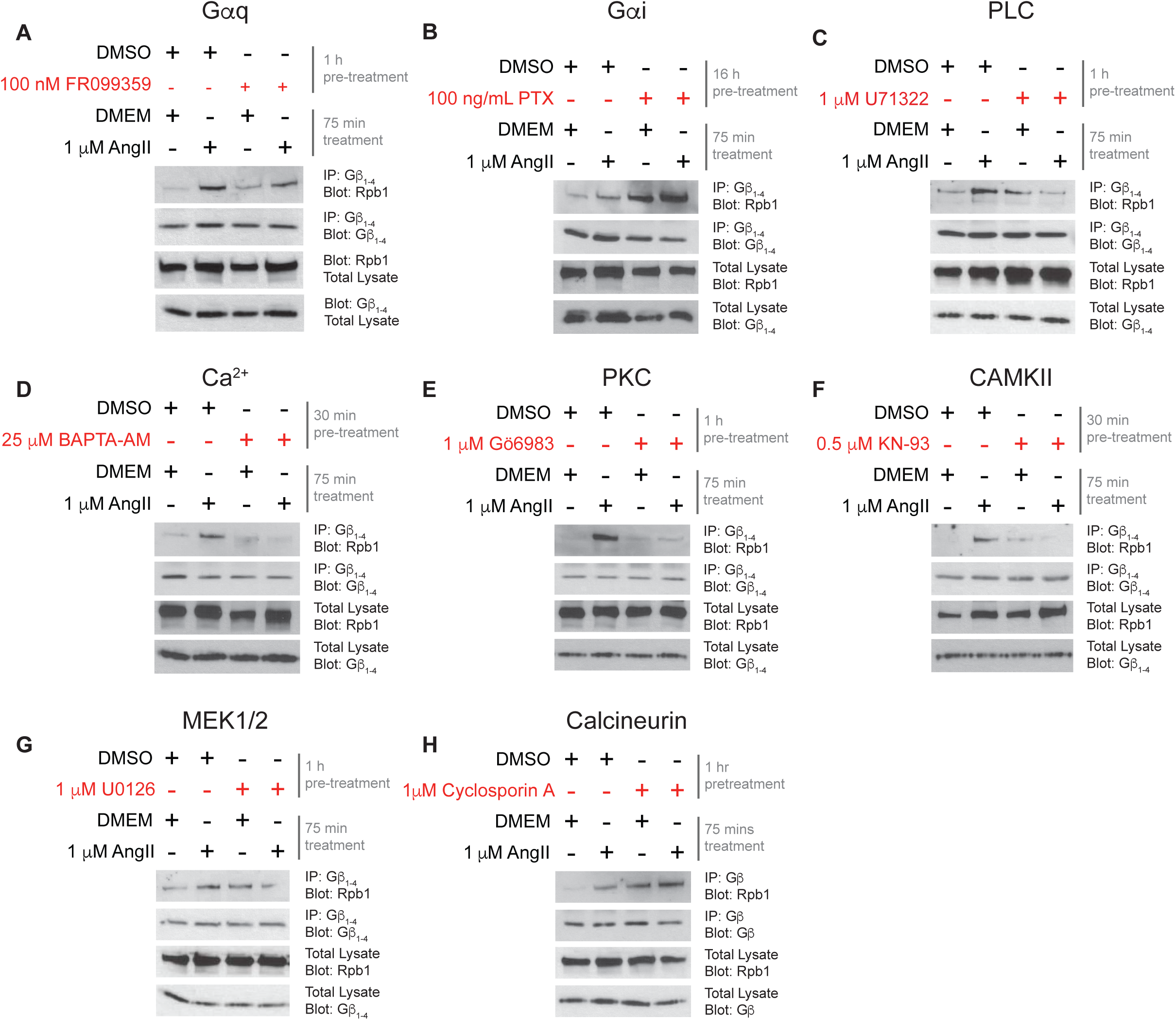
Mechanistic analysis of Gβγ interactions with Rpb1 in rat neonatal cardiac fibroblasts. **(A-H)** Assessment of the effect of inhibition of signalling molecules and effectors implicated in AT1R signalling on the induction of the Gβγ-RNAPII interaction in cardiac fibroblasts. Concentrations of inhibitors and lengths of pre-treatment are indicated in each panel. In all experiments, Ang II treatment was applied at a concentration of 1 µM for 75 min in order to induce the interaction. Data shown is representative of between 3 and 6 independent co-immunoprecipitation and western blot experiments. Corresponding quantification analyses of inhibitor co-IP experiments are depicted in Supplemental Figure 3.

**Figure 3.**
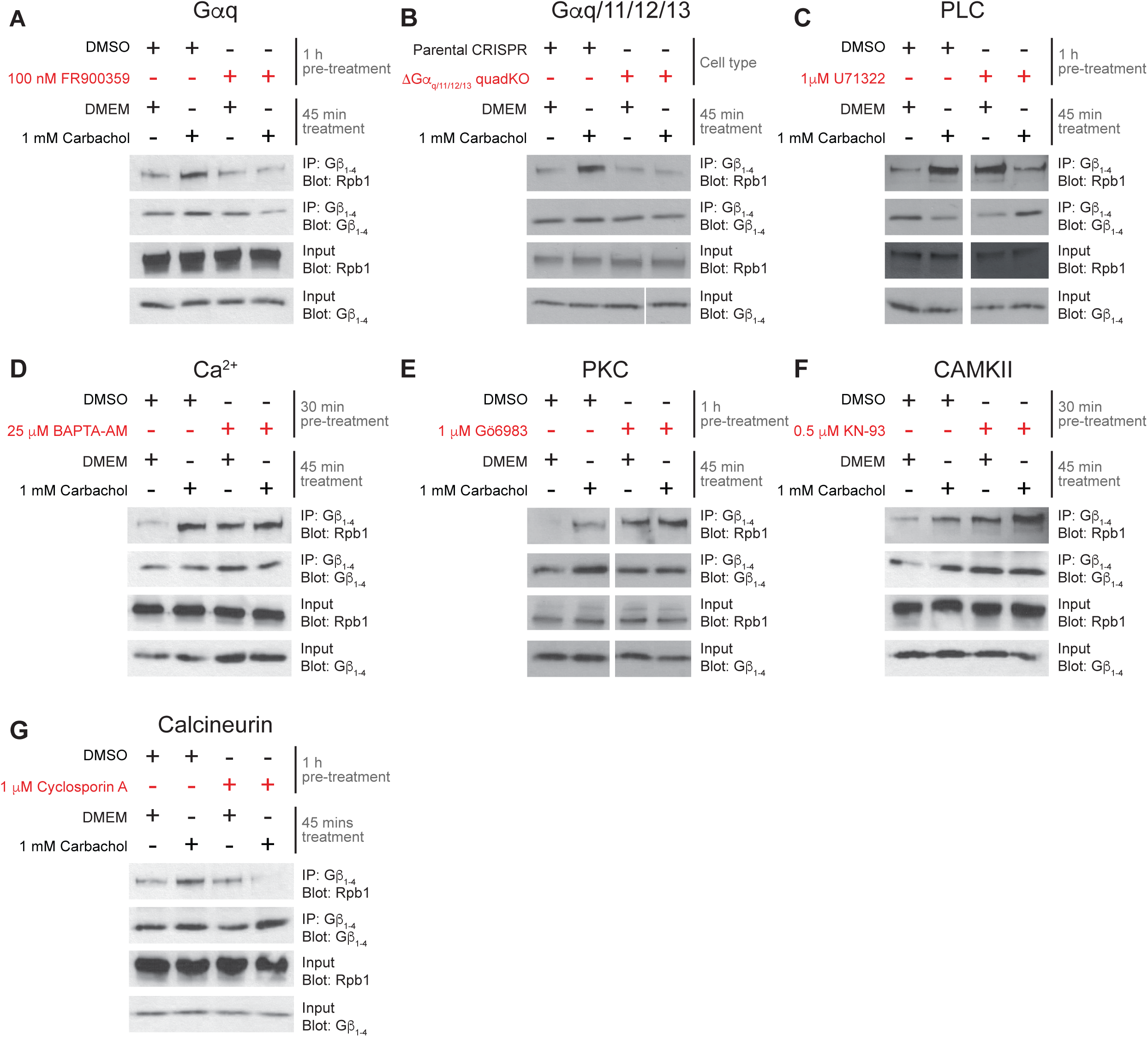
Mechanistic analysis of carbachol-induced Gβγ interaction occurs in HEK 293 cells. **(A-G)** HEK 293 cells were starved for 10-12 hours in DMEM without FBS and were then pre-treated with the indicated inhibitors for the indicated times. Cells were subsequently treated with 1 mM carbachol for 45 min, and the amount of Rpb1 co-immunoprecipitated with Gβ_1-4_ was assessed by western blot. Data is representative of at least 3 independent experiments. The associated quantifications of the co-IPs are represented in Supplemental Figure 4.

Extending these studies to HEK 293F cells, we observed a similar reliance on Gαq signalling for the agonist-induced Gβγ-RNAPII interaction. The carbachol-induced Gβγ-RNAPII interaction was prevented by pre-treatment with the Gαq inhibitor FR900359 (**Figure 3A** **and** Supplemental Figure 4A) and also by CRISPR/Cas9-mediated knockout of Gαq/11/12/13 (**Figure 3B** **and** Supplemental Figure 4B). However, except for this common event, the signalling pathways in cardiac fibroblasts and HEK 293F cells diverged substantially. In HEK 293F cells, U71322 also blocked the carbachol-induced Gβγ-RNAPII interaction but there was a pronounced increase in the basal interaction (**Figure 3C**, Supplemental Figure 4C). Further differences were observed following chelation of calcium with BAPTA-AM which increased basal levels of the Gβγ-RNAPII interaction but did not block further carbachol-induced stimulation of the interaction (**Figure 3D**, Supplemental Figure 4D), suggesting a modulatory role for calcium in HEK 293F cells rather than the direct role seen in cardiac fibroblasts. HEK 293F cells employed different regulatory mechanisms involving protein kinases activated downstream of Gαq/11-coupled GPCRs compared to cardiac fibroblasts. For example, the PKC inhibitor Gö6983 and the CaMKII inhibitor KN-93 both increased basal levels of interaction but did not block carbachol-induced interactions between Gβγ and Rpb1 (**Figure 3E, F**, Supplemental Figure 4E, F). Indeed, inhibition of calcineurin with cyclosporin A blocked the carbachol-mediated increase in interaction between Gβγ and Rpb1, suggesting a role for this phosphatase in mediating the interaction in response to M3-mAChR activation (**Figure 3G** and Supplemental Figure 4G). While the requirement for activation of Gαq is common for the Gβγ-RNAPII interaction in both cell types, the regulation by downstream signalling pathways diverges.

### Roles of individual Gβ subunits in regulating the angiotensin II-activated fibrotic response in rat neonatal cardiac fibroblasts

The Gβ family is comprised of five members which, with the exception of Gβ_5_, exhibit high levels of sequence and structural similarity (Khan et al., 2013). Despite these similarities, Gβ isoforms differ considerably with respect to their associated receptors and signalling pathways (Khan et al., 2015, Yim et al., 2019, Greenwood and Stott, 2019). As our above-reported characterization used a pan-Gβ_1-4_ antibody, we next sought to examine the specificity of Gβ isoforms interacting with Rpb1 in cardiac fibroblasts. We initially focused on Gβ_1_ and Gβ_2_ as they exhibit the highest expression in cardiac fibroblasts determined by RNA-seq (Shu et al., 2018) and RT-qPCR (Supplemental Figure 5A). Immunoprecipitation with a Gβ_1_ specific antibody revealed an increase in the amount of Rpb1 co-immunoprecipitated in response to Ang II treatment, whereas immunoprecipitation of Gβ_2_ indicated a basal interaction with Rpb1 that was lost in response to Ang II treatment (Supplemental Figure 5B). We also assessed Gβ isoform specificity in HEK 293F cells through heterologous expression of FLAG-tagged versions of each Gβ subunit. In response to M3-mAChR activation, FLAG-Gβ_1_ was the only isoform that showed an increased interaction with Rpb1 (Supplemental Figure 5C, D). Hence, an increased interaction between Gβ_1_ and Rpb1 was seen in both cell types, suggesting that our earlier observations using the pan-Gβ_1-4_ antibody likely reflected increased interactions with Gβ_1_.

As we observed isoform-specific roles in RNAPII interactions, we next assessed how knockdown of either Gβ isoform affected the interaction. We first validated knockdown conditions for each Gβ subunit by siRNA at the mRNA and protein levels (Supplemental Figure 6A, B). We observed a reduction in the Ang II-induced Gβγ-RNAPII interaction upon knockdown of Gβ_1_, supporting Gβ_1_ as the isoform involved in the increased interaction with Rpb1. Surprisingly, knockdown of Gβ_2_ also prevented the Ang II-mediated increase in the Gβγ-RNAPII interaction (**Figure 4A, B**). The loss of Gβγ-RNAPII interaction after Gβ_2_ knockdown, despite it not being involved in the Ang II-dependent increase, suggested that AT1R signalling could be altered by loss of Gβ_2_ subunits.

**Figure 4.**
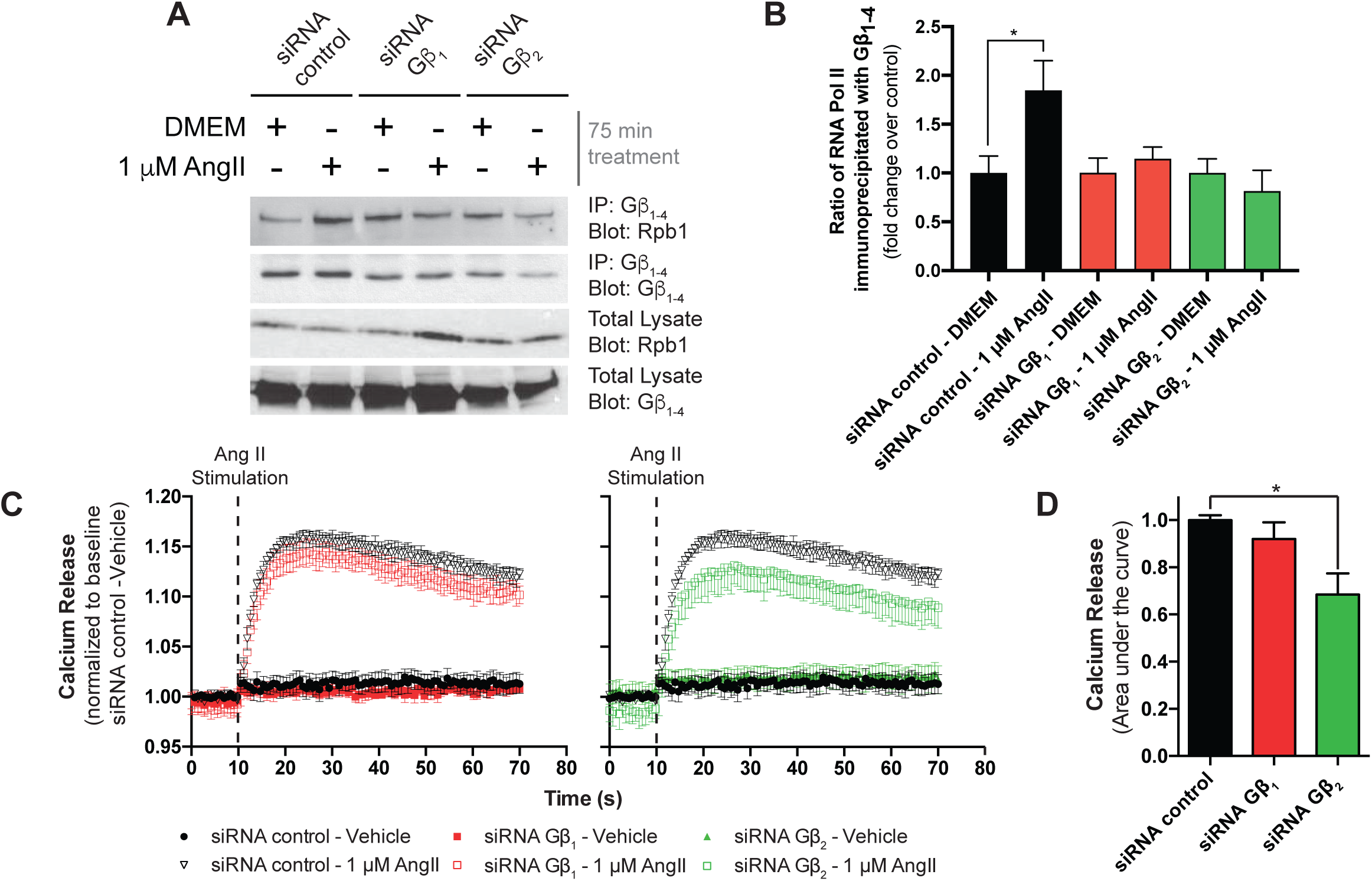
Gβ subunit-specific effects on Ang II signalling and induction of Rpb1 interaction. **(A)** Assessment of the effect of Gβ subunit knockdown by siRNA on the Gβγ-RNAPII interaction upon AT1R stimulation. Cardiac fibroblasts were transfected with siRNA control or siRNA to knockdown Gβ_1_ or Gβ_2_ and were then serum-deprived overnight before treatment with Ang II for 75 min. Cells were assessed for Gβγ-RNAPII interaction by co-immunoprecipitation and western blots. Data represents mean ± S.E.M. of 6-7 independent experiments. **(B)** Densitometry-based quantification of knockdown experiments in (C) were normalized as fold change over the respective siRNA-DMEM condition; data represents mean ± S.E.M. of six independent experiments. **(C)** Traces of calcium release upon AT1R stimulation with Ang II at the 10 s time point, with or without knockdown of either Gβ_1_ or Gβ_2_. Data represents mean ± S.E.M. of fluorescence ratios of 340/516 emission readings to 360/516 emissions readings normalized to basal ratios of three independent experiments. **(D)** Area under the curve analysis of the data obtained in panel A. * indicates p<0.05.

We thus determined whether specific Gβ isoforms were required to initiate signalling cascades proximal to AT1R activation. Following receptor activation, Gβγ subunits regulate intracellular Ca^2+^ mobilization through activation of PLCβ (Park et al., 1993). As we have previously demonstrated Gβ isoform specificity for PLCβ signalling in HEK 293F cells (Khan et al., 2015), we assessed the relative roles of Gβ_1_ and Gβ_2_ in AT1R-dependent Ca^2+^ mobilization. To assess AT1R-dependent intracellular Ca^2+^ mobilization, we used the cell-permeable Ca^2+^ dye Fura 2-AM. Following AT1R activation, we observed a rapid increase in intracellular Ca^2+^ mobilization (**Figure 4A**, black, empty triangles) and the quantified area under the curve (**Figure 4B**, black bar). Knockdown of Gβ_1_ did not alter Ca^2+^ mobilization following stimulation with Ang II (8.1 ± 7.0% decrease, red bar). However, knockdown of Gβ_2_ resulted in a significant 31.6 ± 9% decrease in Ca^2+^ release (**Figure 4A, B**, green bar), suggesting a role for Gβ_2_-containing Gβγ dimers in mediating receptor-proximal signalling downstream of AT1R activation. This suggests Gβ2 knockdown prevented the Ang II-dependent increase in Gβγ-RNAPII interaction through disruption to AT1R Ca^2+^ signalling, aligning with the observed effect of Ca^2+^ chelation with BAPTA-AM. These results highlight the complex interplay between cell surface receptors and multiple Gβγ subunits, in modulating both basal and ligand stimulated RNAPII/Gβγ interactions.

### Gβγ interacts with transcribing RNAPII

As we demonstrated that Gβγ is recruited to RNAPII following AT1R activation, which also activates a transcriptional program in fibroblasts, we assessed the relationship between the transcriptional response and Gβγ recruitment (Shu et al., 2018, Dang et al., 2015). To assess this potential relationship, we disrupted the transcription cycle at two different regulatory points through inhibition of Cdk7, a component of the general transcription factor TFIIH, and Cdk9, the protein kinase subunit of P-TEFb (Zhou et al., 2012). Following RNAPII recruitment, Cdk7 activity stimulates promoter clearance of RNAPII to begin transcription. Soon after RNAPII pauses at a promoter-proximal region and requires the activity of Cdk9 in order to be released into productive elongation (Liu et al., 2015). We assessed involvement of both Cdk7 and Cdk9 on the Ang II-induced Gβγ-RNAPII interaction using the selective inhibitors THZ1 and iCdk9, respectively (Lu et al., 2015, Kwiatkowski et al., 2014). THZ1 abrogated the Ang II-stimulated Gβγ-RNAPII interaction (**Figure 5A**, Supplemental Figure 7A) while iCdk9 resulted in a loss of both the basal and Ang II-stimulated Gβγ-RNAPII interaction (**Figure 5B**, Supplemental Figure 7B). This suggests that the Gβγ-RNAPII interaction requires the transcriptional response to Ang II in cardiac fibroblasts. As with cardiac fibroblasts, in HEK 293F cells disruption of the transcriptional cycle through inhibition of Cdk7 and Cdk9 with DRB also blocked the increased interaction between RNAPII and Gβγ (**data not shown**), showing that the Gβγ/RNAPII interaction is dependent on an active transcriptional response in both cell types.

**Figure 5.**
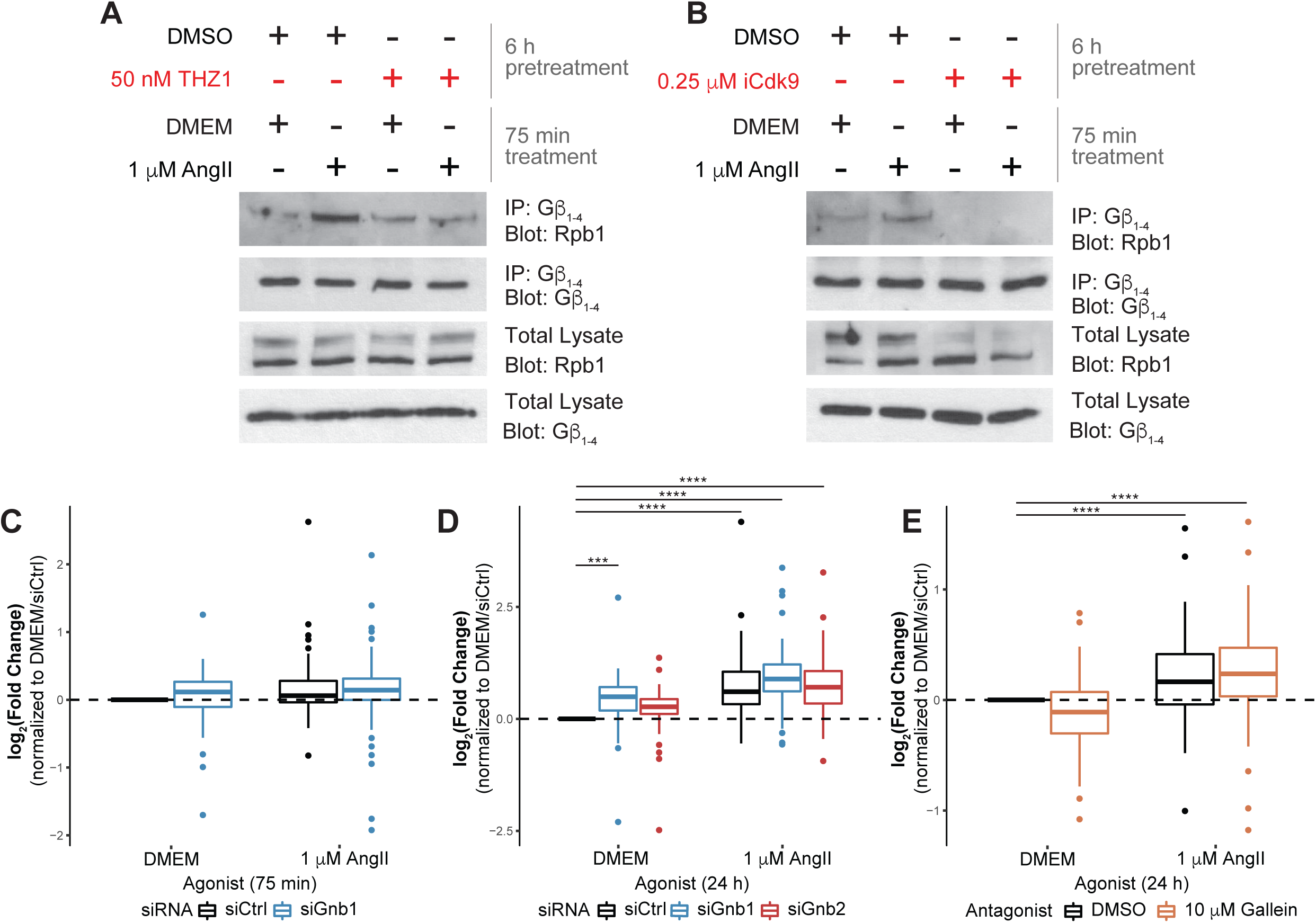
Requirement of RNAPII transcription for Gβγ-RNAPII interaction in rat neonatal cardiac fibroblasts. Effect of Cdk7 inhibition with THZ1 **(A)** or Cdk9 inhibition with iCdk9 **(B)** on Ang II-induced Gβγ-RNAPII interaction. Length of inhibitor pre-treatment is indicated in each respective panel, and the extent of Gβγ-RNAPII interaction was assessed by co-immunoprecipitation coupled to western blot analysis. Data is representative of three independent experiments. Corresponding quantification analyses of inhibitor co-immunoprecipitation experiments are depicted in Supplemental Figure 7. Cumulative log_2_(Fold Change) of all genes detected by qPCR-based fibrosis array following treatment with 1 µM Ang II lasting either 75 min **(C)** or 24 h **(D and E)**. Cardiac fibroblasts were transfected with 50 nM of the indicated siRNA, and were serum-deprived for 12 h before Ang II treatment for the indicated times. Cardiac fibroblasts were pre-treated for 30 min with 10 µM gallein prior to Ang II. Ct values were normalized to the housekeeping genes Hprt1 and Ldha and the log_2_(fold change) over control was determined. For each gene, the average log_2_(fold change) across three independent experiments was plotted. *** indicates p<0.001, **** indicates p<0.0001.

### The role of Gβγ subunits in fibrotic gene expression

In order to understand the role of Gβγ in Ang II-regulated gene expression, we examined changes in the levels of 84 genes involved in the fibrotic response using the Qiagen RT^2^ Profiler^TM^ PCR array platform. Gene expression changes were assessed following 75 min or 24 h Ang II treatment alongside Gβ1 or Gβ2 knockdown. These two time points were selected to investigate the effect of disrupting the Gβγ-RNAPII interaction or, in the longer term, upstream signalling, respectively. We assessed gene expression changes across all 67 genes remaining after excluding genes below our chosen threshold of detection (i.e. Ct > 35). After 75 min of Ang II treatment, we observed a similar upregulation of fibrotic genes in both control and Gβ_1_ knockdown conditions (**Figure 5C**, Supplemental Table 1). However, Gβ_1_ knockdown increased both basal expression and the total number of genes altered by AT1R stimulation (**Figure 5C**, Supplemental Table 1). Following 24 h Ang II treatment, this effect became more pronounced. Gβ_1_ knockdown led to increases in basal gene expression, expression regulated by Ang II treatment and the overall number of genes upregulated (**Figure 5D**, Supplemental Table 1). The increased expression following Gβ_1_ knockdown suggests the Gβγ-RNAPII interaction negatively modulates the Ang II transcriptional response.

Whereas Gβ1 knockdown altered the transcriptional response to Ang II treatment, disruption of AT1R signalling by Gβ_2_ knockdown did not significantly alter basal fibrotic gene expression or the overall response to 24 h Ang II treatment (**Figure 5D**, Supplemental Table 1). The lack of effect of Gβ_2_ knockdown suggests that Gβγ signalling through Ca^2+^ is not required for AT1R-mediated transcriptional changes. To further address the role of Gβγ signalling, we utilized the small-molecule pan-Gβγ inhibitor gallein (Lehmann et al., 2008). As with Gβ_2_ knockdown, pre-treatment with gallein did not significantly alter the transcriptional response following 24 h Ang II treatment (**Figure 5E**). This suggests that Gβγ-dependent signalling downstream of the AT1R is not a key driver of transcriptional changes. Instead, Gβγ is required to modulate processes driven by other signalling pathways and dampen the fibrotic response until such signals rise above a threshold.

### Genome-wide recruitment of Gβ_1_ and the effect on RNAPII occupancy following Ang II treatment

To assess the possibility of genome-wide Gβ_1_ recruitment and changes in RNAPII occupancy following 75 min Ang II treatment in cardiac fibroblasts, we performed chromatin immunoprecipitation followed by next generation sequencing (ChIP-seq) for heterologously expressed FLAG-Gβ_1_ and endogenous Rpb1. We confirmed that, like endogenous Gβ_1_, the interaction of Rbp1 with heterologously expressed FLAG-Gβ_1_ increased following AT1R activation (Supplemental Figure 8A, B). We focused on genes with RNAPII peaks identified by the peak calling software macs2 and annotated with HOMER (Heinz et al., 2010, Zhang et al., 2008). The same Gβ_1_ knockdown conditions that increased the number of genes upregulated in response to Ang II (above) also increased the number of genes occupied by RNAPII following Ang II treatment (**Figure 6A**). To identify groups of genes with similar FLAG-Gβ_1_ and RNAPII occupancy patterns, we performed K-means clustering with genes that RNAPII peaks were identified in any treatment condition. Two K-means clusters were identified (98 genes in cluster 1 and 806 in cluster 2) with distinct occupancy patterns (**Figure 6B, C**). In cluster 1, FLAG-Gβ_1_ occupancy increased within the gene body in response to Ang II. A similar but weaker tendency was also observed in cluster 2 (**Figure 6B**). The increased FLAG-Gβ1 occupancy in cluster 1 corresponded to Gβ1-dependent changes to the Ang II-induced RNAPII occupancy alterations. First, Ang II treatment led to increased RNAPII occupancy throughout the gene body under siRNA control conditions (**Figure 6C**). In the absence of Ang II, Gβ_1_ knockdown increased RNAPII occupancy near transcription start sites (TSSs) which corresponds with increased gene expression under these conditions (**Figure 6C**). Lastly, there was greater RNAPII occupancy when Ang II treatment was combined with Gβ1 knockdown than in the absence of knockdown (**Figure 6C**). Similar RNAPII occupancy patterns were observed in cluster 2, suggesting that Gβ1 also plays a regulatory role along these genes and our FLAG-Gβ_1_ ChIP-seq was not sensitive enough to reliably detect Gβ_1_. We also assessed the functional pathways enriched in cluster 1, through gene ontology (GO) term enrichment. The top four significant GO terms identified (corresponding to cellular processes such as inflammation, fibroblast activation and apoptosis) indicate that Gβ_1_ is recruited to genes involved in processes essential to fibrosis (**Figure 6D**).

**Figure 6.**
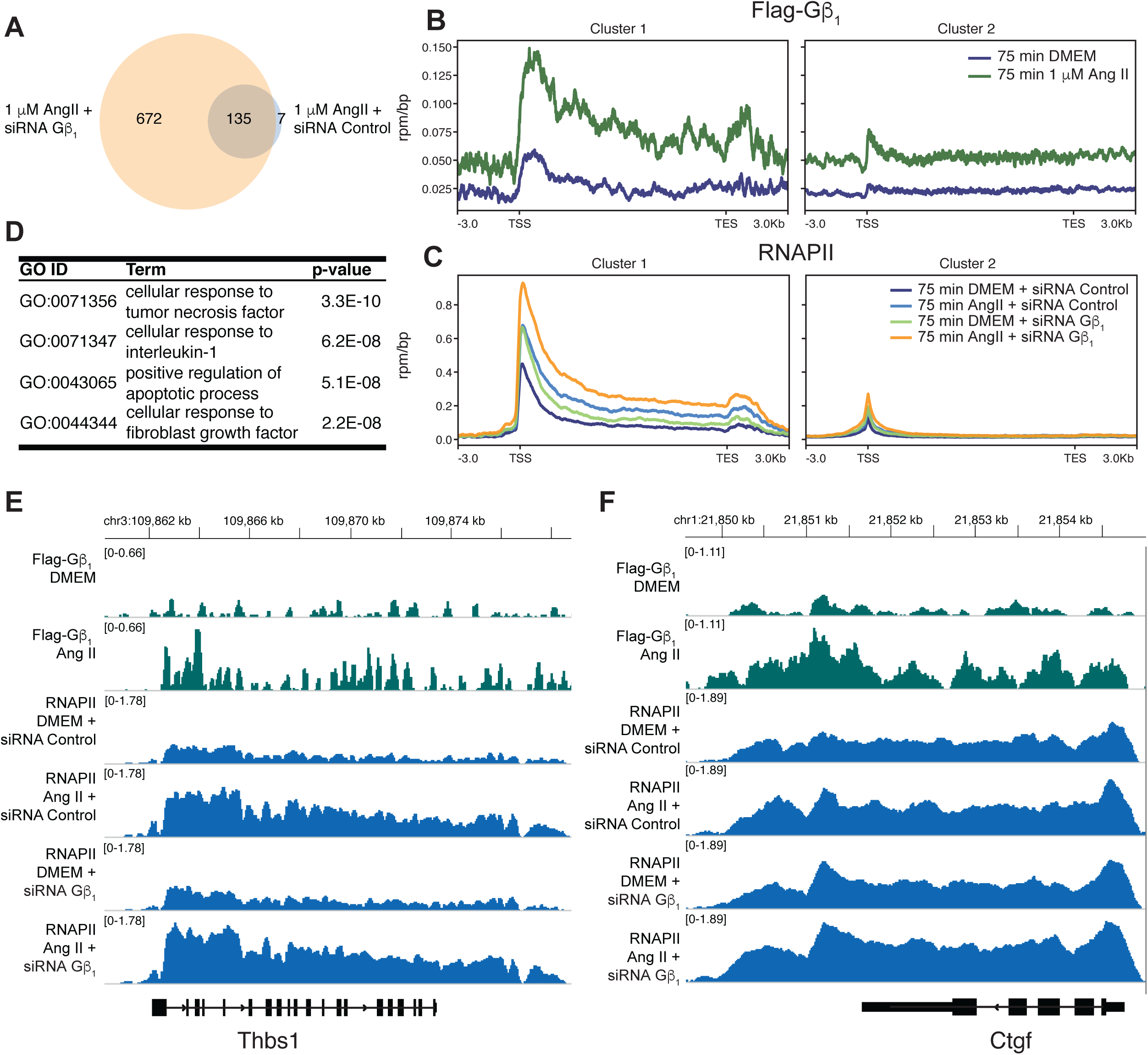
ChIP-seq for FLAG-Gβ1 and Rpb1 following 75 min Ang II treatment in cardiac fibroblasts. Cardiac fibroblasts were transduced with AAV1-FLAG-Gβ_1_ or transfected with the indicated siRNA followed by Ang II treatment (1 µM for 75 min). **(A)** Comparison of genes with annotated RNAPII peaks following Ang II treatment and siRNA control or Gβ_1_. FLAG-Gβ_1_ (**B)** or the Rpb1 subunit of RNAPII **(C)** were immunoprecipitated from crosslinked and sonicated chromatin, followed by DNA purification and next-generation sequencing. Reads were normalized to an exogenous *S. pombe* chromatin spike-in. Genes with a RNAPII peak annotated by HOMER in two of the three replicates were used to identify two K-means clusters. **(D)** Top four significant GO terms enriched in cluster 1. Individual FLAG-Gβ_1_ or RNAPII tracks for two genes from cluster 1, **(E)** Thbs1 or **(F)** Ctgf.

The increased number of genes with RNAPII occupancy in the Ang II and Gβ1 knockdown condition suggested that Gβ_1_ occupancy impairs RNAPII recruitment. As such, we would expect cluster 1 genes to be more enriched in genes with RNAPII occupancy under Ang II and Gβ1 knockdown condition than Ang II and siRNA control conditions. Therefore, we performed a Fisher’s exact test to compare the proportion of cluster 1 genes in these treatment conditions, which demonstrated a significant (p-value < 0.01) enrichment in the Ang II and Gβ1 knockdown condition gene list compared to Ang II and siRNA control condition. This again suggests Gβ_1_ functions to suppress RNAPII transcription following AT1R activation.

In order to assess the relationship between Gβ_1_ occupancy and transcription, we focused on genes from our fibrosis qPCR array that were also found in cluster 1. Eight genes from the fibrosis array were identified in cluster 1, which included five of the seven genes upregulated after 75 min of Ang II treatment such as thrombospondin 1 (Thbs1) and connective tissue growth factor (Ctgf) (**Figure 6E, F**). We confirmed the Ang II-dependent increase in Gβ_1_ occupancy along Ctgf by ChIP-qPCR (Supplemental Figure 8C). We also assessed the effect of Gβ_1_ knockdown on AT1R-dependent changes of RNAPII occupancy along Ctgf by ChIP-qPCR. Similar to our ChIP-seq analysis, we observed a greater increase in RNAPII along the gene in response to Ang II under siRNA GNB1 knockdown compared to siRNA control, where we observed a slight decrease (Supplemental Figure 8D). We also validated the change in expression of Ctgf by RT-qPCR using primers designed in-house (Supplemental Table 2). Under control conditions Ang II had a minor effect on Ctgf expression, however in the absence of Gβ_1_ Ang II treatment resulted in a significant upregulation of Ctgf mRNA. (Supplemental Figure 8E). Taken together, our results demonstrate Gβ_1_ recruitment negatively regulates expression of genes involved in the fibrotic response to Ang II by inhibiting early stages of the RNAPII transcription cycle.

## DISCUSSION

The functional specificity of Gβ and Gγ subunits has been mostly investigated in the context of signalling proximal to GPCR activation (i.e., the regulation of effector activity downstream of receptor stimulation) (Khan et al., 2013). In contrast, our findings provide new insights regarding non-canonical roles of specific Gβγ dimers in more distal events in the nucleus, particularly in the regulation of gene expression. Here, we demonstrate for the first time an interaction between Gβγ and RNAPII and investigate the regulatory signalling mechanisms in transformed cell lines (HEK 293 cells) and in primary cells (neonatal rat cardiac fibroblasts). The interaction of Gβγ and RNAPII represents a significant addition to the expanding list of Gβγ interactors, and our findings suggest that regulatory mechanisms impacting the interaction are dependent on cellular context. We also show that Gβγ signalling is a critical regulator of the fibrotic response in cardiac fibroblasts.

Our findings suggest that following acute treatment with Ang II, Gβ_1_ is transiently recruited to pro-fibrotic genes to negatively regulate RNAPII recruitment, thereby limiting the fibrotic response following transient fluctuations in local Ang II concentrations likely seen *in vivo*. This negative RNAPII regulation may potentially occur through direct interactions with RNAPII, preventing its recruitment or other aspects of initiation, or else via an indirect mechanism in which Gβγ would form part of a larger RNAPII-containing complex altering the local chromatin landscape. We cannot currently distinguish between these two possibilities, given that our co-immunoprecipitation assay was performed using whole-cell lysates. On the other hand, chronic stress or damage to the heart leads to a sustained increase of Ang II concentrations in cardiac tissue (Sun and Weber, 1996, Passier et al., 1996). We propose that such sustained AT1R signalling overcomes the transient Gβ_1_ “brake” to elicit a robust fibrotic response. Alternatively, pro-fibrotic factors that are upregulated and secreted following AT1R activation may elicit autocrine signalling pathways that overcome the Gβ_1_ transcriptional repression (Lee et al., 1995, Ma et al., 2018). Our gene expression data at 75 min, and more especially at 24 h, begins to identify the increased number and greater gene expression in the absence of the proposed negative regulatory mechanism when Gβ_1_ is knocked down. Further analysis of the kinetics of the interaction and how this changes the dynamics of chromatin occupation or gene expression are required as well. Future experiments assessing nascent RNA production are required to accurately determine gene expression changes at early time points.

We demonstrated that Gβ_2_, and not Gβ_1_, was important for proximal signalling downstream of AT1R activation similar to the requirement of specific Gβ isoforms for activation of PLCβ in HEK 293 cells (Khan et al., 2015). Our data suggest that Gβ_2_ plays a minimal role in regulating AT1R-dependent gene expression *per se*. Rather, our findings using the broad-spectrum Gβγ inhibitor gallein suggest that receptor-proximal Gβγ signalling in general is not required for the transcriptional response and instead it is dependent on Gαq signalling and more distal Gβ_1_-dependent events. Knockdown of Gβ_2_ also compromised Ang II-mediated interactions between Gβγ and RNAPII even though Gβ_2_ had a limited role in the fibrotic transcriptional response. This suggests Gβ_2_ knockdown does not prevent the response but rather alters the kinetics of Gβγ-RNAPII interactions, which then translates into different fibrotic responses over time. Further, the roles of specific Gγ subunits in mediating proximal signal transduction must also be considered as for other Gβγ effectors (Khan et al., 2015), and should be the subject of future studies. Taken together, our findings suggest that in fibrosis and potentially in other diseases, the indiscriminate targeting of Gβγ signalling (e.g. with compounds such as gallein) will result in outcomes that differ considerably from those obtained by targeting particular Gβγ combinations (Lin and Smrcka, 2011, Kamal et al., 2011, Smrcka et al., 2008).

Analysis of the signalling networks regulating the Gβγ/RNAPII interaction yielded four main conclusions: (1) different GPCR signalling systems in distinct cell types lead to different kinetics of the Gβγ-RNAPII interaction, (2) different signalling pathways downstream of GPCR activation act to both induce or modulate the interaction, (3) Gαq-coupled GPCRs regulate the interaction in both cell types examined, and (4) signalling ultimately converged on activation of transcription. Indeed, our results suggest that the cell context plays a critical role in determining the mechanism by which the Gβγ-RNAPII interaction is regulated. First, in cardiac fibroblasts, the Gβγ/RNAPII interaction depended on a Gq-PLCβ-Ca^2+^-CaMKII/PKC/MEK-dependent pathway downstream of AT1R activation, whereas calcineurin acted as a basal negative regulator (summarized in Supplemental Figure 9). On the other hand, in HEK 293 cells, we observed that the interaction was reliant on a Gq-PLCβ-Ca^2+^-calcineurin pathway downstream of M3-mAChR activation, whereby PKC and CaMKII both negatively regulate this interaction under basal conditions (summarized in Supplemental Figure 9). The involvement of Ca^2+^, PKC and ERK1/2 in the induction of the Gβγ/RNAPII interaction in fibroblasts is supported by previous reports that demonstrate their involvement in Ang II-induced fibrosis (Chintalgattu and Katwa, 2009, Olson et al., 2008).

The different signalling pathways promoting the Gβγ-RNAPII interaction appear to converge at the point of Cdk7 and Cdk9 activation. In particular, we found that the Cdk7 and Cdk9 inhibitors (DRB, THZ1 and iCdk9, respectively) inhibited both carbachol-induced Gβγ-RNAPII interaction in HEK 293 cells and the analogous Ang II-induced interaction in cardiac fibroblasts. This suggests the differential regulatory signalling pathways identified are due to cell type- and receptor-specific activation pathways of both Cdk7 and Cdk9. The recruitment of Gβγ serves as a common negative regulatory mechanism regardless of the pathway leading to transcriptional activation. Furthermore, a strong connection has been established between the control of transcriptional pausing and pathological cardiac remodelling, although primarily in the cardiomyocyte (Yang et al., 2017, Sayed et al., 2013, Anand et al., 2013, Duan et al., 2017, Stratton et al., 2016, Sano et al., 2002). Our results indicate that regulation of the early stages of the RNAPII transcription cycle is also an important checkpoint in the fibrotic response mediated by cardiac fibroblasts.

Taken together, the Gβγ-RNAPII interaction identifies a new mechanism by which Gβγ modulates gene expression. Our study highlights the complex interplay of different Gβγ subunit combinations at the cell surface and in the nucleus initiated upon stimulation of Gαq-coupled receptors. Since Gβ_1_γ dimers play an important role in regulating the expression of fibrotic genes in cardiac fibroblasts, the development of selective Gβ_1_γ inhibitors hold some promise for preventing the pathological consequences of myocardial damage.

## METHODS

### Reagents

The following were all purchased from Sigma-Aldrich: carbachol, angiotensin II, BAPTA-AM, KN-93, Gö6983, PTX, U0126, calyculin A, cyclosporin A, TRI reagent, isopropyl thiogalactopyranoside (IPTG), protease inhibitor cocktail, triton X-100, bovine serum albumin, ethylenediaminetetraacetic acid (EDTA), 70% NP-40 (Tergitol), sodium deoxycholate, magnesium chloride, lithium chloride, anti-rabbit IgG (whole molecule)-agarose antibody, anti-mouse IgG (whole molecule)-agarose antibody, goat anti-rabbit IgG (whole molecule) conjugated to peroxidase secondary antibody, goat anti-mouse IgG (Fab specific) conjugated to peroxidase secondary antibody, anti-FLAG M2 antibody, and rabbit IgG (St. Louis, MO, USA). U71322 pan-PKC inhibitor was purchased from Biomol International (Plymouth Meeting, PA, USA). Lysozyme (from hen egg white) and phenylmethylsulfonyl fluoride (PMSF) were purchased from Roche Applied Sciences (Laval, QC, Canada). Ethylene glycol bis (2-aminooethyl ether) N,N,N’,N’ tetraacetic acid (EGTA) and HEPES were purchased from BioShop (Burlington, ON, Canada). Sodium chloride, glutathione (reduced form), dithiothreitol (DTT) and Dynabeads protein G were purchased from Fisher Scientific (Ottawa, ON, Canada). Dulbecco’s modified Eagle’s medium (DMEM) (supplemented with 4.5 g/L glucose, L-glutamine and phenol red), DMEM low glucose (supplemented with 1.0 g/L glucose, L-glutamine and phenol red), Hank’s Balanced salt solution (HBSS), HBSS (with no phenol), Penicillin/Streptomycin solution, Tris base buffer, ampicillin sodium salt, and fetal bovine serum were purchased from Wisent (St. Bruno, QC, Canada). Glutathione sepharose 4B GST beads was purchased from GE Healthcare (Mississauga, ON, Canada). Lipofectamine 2000 and Alexa Fluor 488 goat anti-mouse IgG were purchased from Invitrogen (Burlington, ON, Canada). Enhanced chemiluminescence (ECL) Plus reagent was purchased from Perkin Elmer (Woodbridge, ON, Canada). Moloney murine leukemia virus reverse transcriptase (MMLV-RT) enzyme and recombinant RNasin® ribonuclease inhibitor were purchased from Promega (Madison, WI, USA). Evagreen 2X qPCR MasterMix was purchased from Applied Biological Materials Inc. (Vancouver, BC, Canada) and iQ SYBR Green Supermix was purchased from Bio-Rad Laboratories (Mississauga, ON, Canada). Anti-Gβ1-4 (T-20) antibody, anti-RNA Polymerase I Rpa194 (N-16) antibody, anti-ERK1/2 antibody, anti-Gαq antibody and anti-Rpb1 (N20) were purchased from Santa Cruz Biotechnology, Inc. (Dallas, TX, USA). Anti-RNA polymerase II clone CTD4H8 (Rpb1) antibody was purchased from EMD Millipore (Temecula, CA, USA). Anti-*Schizosaccharomyces pombe* histone H2B (ab188271) antibody was purchased from Abcam Inc. (Toronto, ON, Canada). Polyclonal anti-Gβ_1_ and anti-Gβ_2_ were a generous gift of Professor Ron Taussig (UT Southwestern). THZ1 was a gift from Nathanael S. Gray (Harvard University) and iCdk9 was a gift from James Sutton (Novartis). FLAG-Gβ_1_, FLAG-Gβ_2_, FLAG-Gβ_3_, FLAG-Gβ_4_ and FLAG-Gβ_5_ plasmids were obtained from UMR cDNA Resource (www.cdna.org).

### Tissue culture, transfection and treatments

Human embryonic kidney 293 (HEK 293), HEK 293T cells and CRISPR/Cas9 generated ΔGαq/11/12/13 knockout HEK 293 cells (quadKO cells) (Devost et al., 2017), a generous gift from Dr. Asuka Inoue (Tohuku University, Sendai, Japan), were grown at 37°C in 5% CO_2_ in DMEM supplemented with 5% (v/v) fetal bovine serum and 1% (v/v) penicillin/streptomycin (P/S). HEK 293 cells were transiently transfected with FLAG-Gβ1-5 using Lipofectamine 2000 as per the manufacturer’s recommendations. Primary rat neonatal cardiac fibroblasts were isolated from 1-3 day old Sprague-Dawley rat pups (Charles River Laboratories, St-Constant, Quebec) as previously described (Calderone et al., 1998). All procedures using animals were approved by the McGill University Animal Care Committee, in accordance with Canadian Council on Animal Care Guidelines. Two days after isolation, cells were detached with trypsin/EDTA and plated at a density of ∼8 x 10^3^ cells/cm^2^ in fibroblast growth medium for 48h. For siRNA transfection, cardiac fibroblasts were plated at a density of ∼20 x 10^3^ cells/cm^2^ and transfected using Lipofectamine 2000 as per the manufacturer’s instructions. For treatment of HEK 293F cells, HEK 293F quadKO cells or cardiac fibroblasts, cells were serum-deprived for 6 h with DMEM or overnight (∼12 h) with DMEM low glucose (with no FBS and no P/S) respectively, and subsequently treated with pathway inhibitors, 1 mM carbachol or 1 µM Ang II for the treatment lengths indicated in the various assays.

### RT-qPCR

Reverse transcription of RNA isolated from rat neonatal cardiac fibroblasts was performed as previously described (Khan et al., 2015). Briefly, cells were lysed in TRI reagent and RNA was extracted using a protocol adapted from Ambion (Burlington, ON, Canada). Reverse transcription was performed on 1 µg of total RNA using an MMLV-RT platform according to the manufacturer’s protocol. Subsequent qPCR analysis was performed with Evagreen Dye qPCR master-mixes using a Corbett Rotorgene 6000 thermocycler or Bio-Rad 1000 Series Thermal Cycling CFX96 Optical Reaction module. mRNA expression data were normalized to housekeeping transcripts for U6 snRNA. Ct values obtained were analyzed to calculate fold change over respective control values using the 2^-**ΔΔ**Ct^ method. Primer sequences for all primers used are listed in **Supplemental Table 2**.

### Ca^2+^ mobilization

Cardiac fibroblasts were cultured as previously described following transfection with respective siRNA. Cardiac fibroblasts were washed and media replaced with HBSS (no phenol) and incubated for 1 h at 37°C and 5% CO_2_. Media was replaced with Fura 2-AM in HBSS and incubated for another 1 h at 37°C and 5% CO_2_. Fura 2-AM containing media was replaced with HBSS and Cardiac fibroblasts incubated for another 30 min at 37°C and 5% CO_2_ prior to recordings. Baseline recordings were obtained every 0.7 s for 10 s followed by injection of Ang II to a final concentration of 1 µM and recordings obtained every 0.7 s for a total of 1 min. A control well with no Fura-2 AM was included in order to control for background fluorescence. Fluorescence intensity was recorded using Bio-Tek Synergy 2 Multi-Mode Microplate Reader with fluorescence excitation at 340 nm or 360 nm and fluorescence emission at 516 nm. Data is presented as the ratio of fluorescence emission at 516 nm following 340 nm excitation over 360 nm excitation. The ratio was normalized to the mean baseline ratio from control cells.

### Nuclear isolation

Nuclei from HEK 293 cells and cardiac fibroblasts were isolated as previously described (Campden et al., 2015b). Briefly, cells seeded in T175 flasks (Corning) were treated as indicated, washed three times with 1X PBS (137 mM NaCl, 2.7 mM KCl, 10 mM Na_2_HPO_4_, 1.8 mM KH_2_PO_4_), and harvested in 1X PBS by centrifugation. Pelleted cells were lysed in lysis buffer (320mM sucrose, 10 mM HEPES, 5 mM MgCl_2_, 1 mM DTT, 1 mM PMSF, 1% Triton X-100), added gently on top of a high-sucrose buffer (1.8 M sucrose, 10 mM HEPES, 5 mM MgCl_2_, 1 mM DTT, 1 mM PMSF), and centrifuged at 4600 g for 30 min at 4°C, separating unlysed nuclei from the cytosolic fraction. Pelleted nuclei were then resuspended in resuspension buffer (320 mM sucrose, 10 mM HEPES, 5 mM MgCl_2_, 1 mM DTT, 1 mM PMSF), pelleted at 300 g for 5 min and subsequently lysed in 1X RIPA buffer.

### Immunoprecipitation and western blotting

Immunoprecipitation (IP) assays of Gβ and Rpb1 pull downs were performed as previously described, with minor alterations (Robitaille et al., 2010). Protein extracts from treated HEK 293 cells and cardiac fibroblasts lysed in RIPA (1% NP-40, 50 mM Tris-HCl ph 7.4, 150 mM NaCl, 1 mM EDTA, 1 mM EGTA, 0.1% SDS, 0.5% sodium deoxycholate) were quantified by Bradford assay and 500 µg of protein lysate was precleared with 15 µl of anti-rabbit IgG-agarose beads. Precleared lysates were then incubated with 1 µg anti-Gβ_1-4_, 2 µg of anti-Rpb1 or anti-Gβ_1_ serum or anti-Gβ_2_ serum overnight at 4°C with end-over mixing. The next day, 40 µl of washed agarose beads were added to each lysate/antibody mixture, incubated for 3.5 hours at 4°C with end-over mixing, and then beads were washed 3X with RIPA. Proteins were eluted off the beads by the addition of 4X Laemmli buffer followed by denaturation at 65°C for 15 min. Protein immunoprecipitation and co-IP were then assessed by western blot as previously described (Khan et al., 2015). Resulting western blot images were quantified using ImageJ 1.48v.

### Rat Fibrosis qPCR arrays

Fibrosis qPCR arrays were performed as per the manufacturer’s instructions (Qiagen, Toronto, ON, Canada). Briefly, 0.5 µg of isolated total RNA from siRNA transfected and vehicle or Ang II treated cardiac fibroblasts was subject to genomic DNA elimination using mixes supplied with the array kit for 5 mins at 42°C. DNA eliminated RNA was then subject to reverse transcription reactions using Qiagen RT^2^ First Strand Kits with protocols according to the manufacturer’s instructions. Qiagen RT^2^ SYBR Green MasterMix was added to the cDNA and subsequently dispensed in wells of a 96-well plate containing pre-loaded lyophilized primers provided by the manufacturer. Quantitative PCR reactions were then run on an Applied Biosystems ViiA 7 thermocycler according to the manufacturers cycle recommendations. Each sample was run on separate individual 96 well plates and Ct values for each gene assessed were collected and analyzed; Ct values greater than 35 were eliminated from the overall analysis. Expression data was normalized to levels of two housekeeping genes contained on each plate – Ldha1 and Hprt.

### AAV Production and transduction of cardiac fibroblasts

FLAG-Gβ_1_ and FLAG-Gβ_2_ were PCR amplified from a pcDNA3.1+ plasmid and BamHI and EcoRI restrictions sites added to the 5’ and 3’ end, respectively. These restrictions sites were used to insert each FLAG-Gβ into the pAAV-CAG plasmid. Adeno-associated viruses were produced as previously described (Burger and Nash, 2016). Cells were transduced with AAV1-FLAG-Gβ_1_ (MOI of 10^3^ or 5×10^4^) in DMEM low glucose for 6h. Additional media was added to obtain a final 7% FBS concentration and incubated for another 24 h. At this point, the cells were detached with trypsin/EDTA and plated as described for respective experiments.

### ChIP-qPCR

Immunoprecipitation in cardiac fibroblasts was performed as previously described, with minor modifications (Bolli et al., 2013). Isolated nuclei were sonicated with a Diagenode BioRuptor^TM^ UCD-200 (18 cycles, 30 s on/off, high power) to shear chromatin. FLAG-Gβ_1_ immunoprecipitation was performed with 10 μg sheared rat chromatin alongside 5 μg of *Schizosaccharomyces pombe* yeast chromatin, obtained as previously described (Mbogning and Tanny, 2017). Chromatin was immunoprecipitated with an anti-FLAG M2 antibody (2 µg) or equivalent amount of rabbit IgG alongside an anti-*Schizosaccharomyces pombe* H2B antibody. RNAPII immunoprecipitation was performed with 20 µg of sheared rat chromatin alongside 0.2 µg of *Schizosaccharomyces pombe* yeast chromatin. Chromatin was immunoprecipitated with an anti-Rpb1 (8WG16) antibody. Localization was assessed by qPCR with primers for specific genomic loci (Supplemental Table 2). All qPCR reactions were performed using a Bio-Rad 1000 Series Thermal Cycling CFX96 Optical Reaction module and iQ SYBR Green Supermix. Data analysis included subtracting the % Input of IgG control for each treatment from the respective IP, followed by normalization to the % Input yeast *cdc2^+^* of each FLAG IP or *act1^+^* for each RNAPII IP to account for differences in IP efficiencies.

### ChIP-seq immunoprecipitation and data analysis

Immunoprecipitation in Cardiac fibroblasts was performed as previously described, with minor modifications (Bolli et al., 2013). Isolated nuclei were sonicated with a Diagenode BioRuptor^TM^ UCD-200 (18 cycles, 30 s on/off, high power) to shear chromatin. FLAG-Gβ_1_ immunoprecipitation was performed with 40 μg of sheared rat chromatin alongside 0.4 μg of chromatin from a *S. pombe* strain expressing FLAG-Bdf2. RNAPII immunoprecipitation was performed using 20 µg sheared rat chromatin alongside 0.2 µg wild-type *S. pombe* chromatin. Chromatin was immunoprecipitated with an anti-FLAG M2 antibody (2 µg) or anti-Rpb1 (8WG16) antibody (2 µg). Two biological replicates of FLAG-Gβ_1_ immunoprecipitation and three biological replicates of Rpb1 immunoprecipitation were included. Following immunoprecipitation and DNA cleanup, libraries were prepared with the NEBNext® Ultra^TM^ II DNA Library Prep kit for Illumina and 50 bp single end reads obtained with an Illumina HiSeq 4000 at the McGill University and Génome Québec Innovation Centre, Montréal, Canada.

Reads were trimmed with TrimGalore (0.6.0) (Krueger, Martin, 2011) using the following settings: --phred33 --length 36 -q 5 --stringency 1 -e 0.1. A Bowtie2 genome comprised of the Ensembl rat reference genome (Rattus.norvegicus.Rnor.6.0.94) (Zerbino et al., 2018) and *S. pombe* reference genome (Schizosaccharomyces_pombe.ASM294v2) was built with the bowtie2-build function. Processed reads were aligned to the custom combined rat and *S. pombe* genome with Bowtie2 (v2.3.5), followed by removal of low-quality mapped reads (MAPQ < 10) and reads mapped to non-standard chromosomes with SAMtools (v1.9) (Li et al., 2009). Duplicate reads were removed with Picard tools (v2.20.6, Broad Institute). Aligned reads were separated into individual files for the rat or *S. pombe* genome respectively. For RNAPII ChIP, peaks were called using macs2 (v2.1.1) with settings --broad and --broad-cutoff 0.1 (Zhang et al., 2008), reads extended by the fragment length determined by phantompeakqualtools (v1.14) (Landt et al., 2012, Kharchenko et al., 2008), a scaling factor estimated using the NCIS R package (Liang and Keles, 2012) and potential misassembled regions of the rat genome blacklisted (Ramdas et al., 2019). RNAPII peaks were annotated with HOMER (v4.11) (Heinz et al., 2010) and those protein-coding genes with RNAPII peaks in two of three replicates in any treatment were used for subsequent analysis. BAM files of treatment replicates were combined, input reads subtracted with the deepTools (Ramirez et al., 2014) function bamCompare (--scaleFactorsMethod SES) and negative values set to 0. Lastly, values were converted to counts per million mapped reads with library size adjusted by the total number of reads aligned to the *S. pombe* genome. K-means clustering for genes with identified RNAPII peaks and data visualization was performed with the deepTool’s computeMatrix and plotProfile functions. Gene ontology enrichment was performed using the R package topGO (v2.36.0).

### Statistical Analysis

Statistical tests were performed using GraphPad Prism 8.0 software. For quantifications of immunoprecipitation experiments, two-way analysis of variance (ANOVA) followed by post-hoc Dunnett’s test was used on quantifications of western blot bands, with all multiple comparisons being made to vehicle-vehicle conditions. To analyse Ca^2+^ release experiments, the dependent measure was the area under the curve (AUC), computed from release-time data sets. AUC data were subjected to ANOVA and Dunnett’s tests, using as a point of comparison the siRNA control condition. For the FLAG-Gβ and RNAPII interaction in HEK 293F cells, one-sample t-tests were performed with a Bonferroni correction. Summary gene expression of the fibrosis array qPCR was compared with a two-way ANOVA followed by post-hoc t-tests with a Bonferroni correction. Individual gene expression from the fibrosis qPCR array and Ctgf gene expression with in-house primers was assessed with a two-way ANOVA followed by Bonferroni corrected post-hoc t-tests at individual time points. For validation of Gβ_1_ and Gβ_2_ knockdown in cardiac fibroblasts, fold changes over siRNA control were compared to siRNA control using paired Student’s t-tests. For FLAG-Gβ_1_ ChIP-qPCR, independent paired Student t-tests with a Bonferroni post-hoc correction were performed. A Fisher’s exact test was used to compare the proportion of cluster 1 genes in the list of genes with RNAPII peaks following Ang II treatment in control or Gβ_1_ knockdown conditions and to assess GO Term enrichment in cluster 1 genes. Alpha was set at p<0.05 (2-tailed). All results are expressed as mean ± S.E.M, and data are represented as pooled experiments whose sample sizes are indicated in figure legends.

## Acknowledgements

This work was supported by a grant from the Heart and Stroke Foundation of Canada (G-15-0008938) to T.E.H and J.C.T, a grant from Canadian Institute of Health Science (CIHR) (MOP 130362) to J.C.T. and a grant from CIHR (PJT-159687) to T.E.H. R.M. and S.M.K were supported by studentships from the McGill-CIHR Drug Development Training Program and the McGill Faculty of Medicine. We thank Dr. Ron Taussig (UT Southwestern), Dr. Nathanael S. Gray, and Novartis for providing materials instrumental to this study. Lastly, we thank all the members of the Hébert and Tanny labs for discussion and critical reading of the manuscript.

## Author Contributions

S.M.K, R.D.M, J.C.T, and T.E.H designed the experiments and wrote the manuscript. S.M.K, R.D.M, S.G, C.B, J.J.T, A.Z, S.M, and P.T conducted experiments. S.M.K, R.DM and C.B analyzed experiments. R.D.M, J.J.T, J.C.T and T.E.H edited the manuscript.

---

**Supplemental Figure 1.**
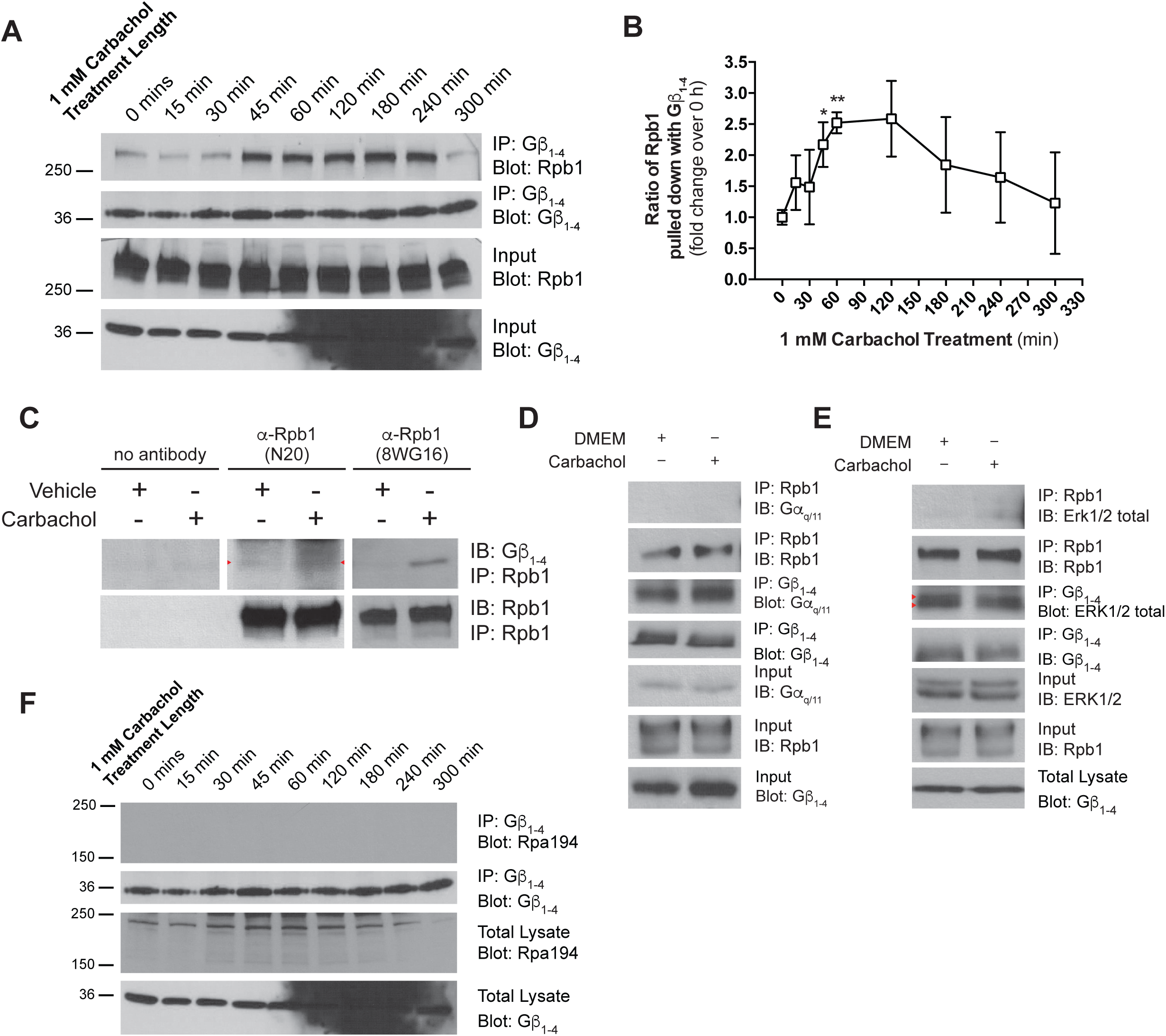
Induction of the Gβγ-RNAPII interaction in HEK 293 cells. **(A)** Time-course analysis of the induction of the Gβγ-RNAPII interaction. The amount of Rpb1 co-immunoprecipitated with Gβ_1-4_ from HEK 293 cells treated for the indicated times with 1 mM carbachol was assessed by western blot for each time point. Data is representative of three independent experiments. **(B)** Quantification of Gβγ-RNAPII time-course co-immunoprecipitation. Densitometry-based analysis of bands corresponding to Rpb1 at each timepoint was normalized to the band intensity of the amount of Gβ_1-4_ immunoprecipitated to yield ratios of Rpb1 pulled down with Gβ_1-4_. **(C)** Assessing the Gβγ and Rpb1 interaction by immunoprecipitation of Rpb1 with two different antibodies. Western blots are representative of at least two independent experiments. Immunoprecipitation experiments demonstrating that carbachol treatment does not induce interaction of Rpb1 with **(D)** Gα_q/11_ nor **(E)** ERK1/2 in HEK 293 cells, and also does not alter the amount of Gα_q/11_ or ERK1/2 interacting with Gβγ under such conditions. **(F)** Assessment of interaction between Gβ_1-4_ and Rpa194, the largest subunit of RNA polymerase I. Data represents analysis of a time course experiment western blot performed as in Supplemental Figure 2A. Data represents mean ± S.E.M; * indicates p<0.05, ** indicates p<0.01.

**Supplemental Figure 2.**
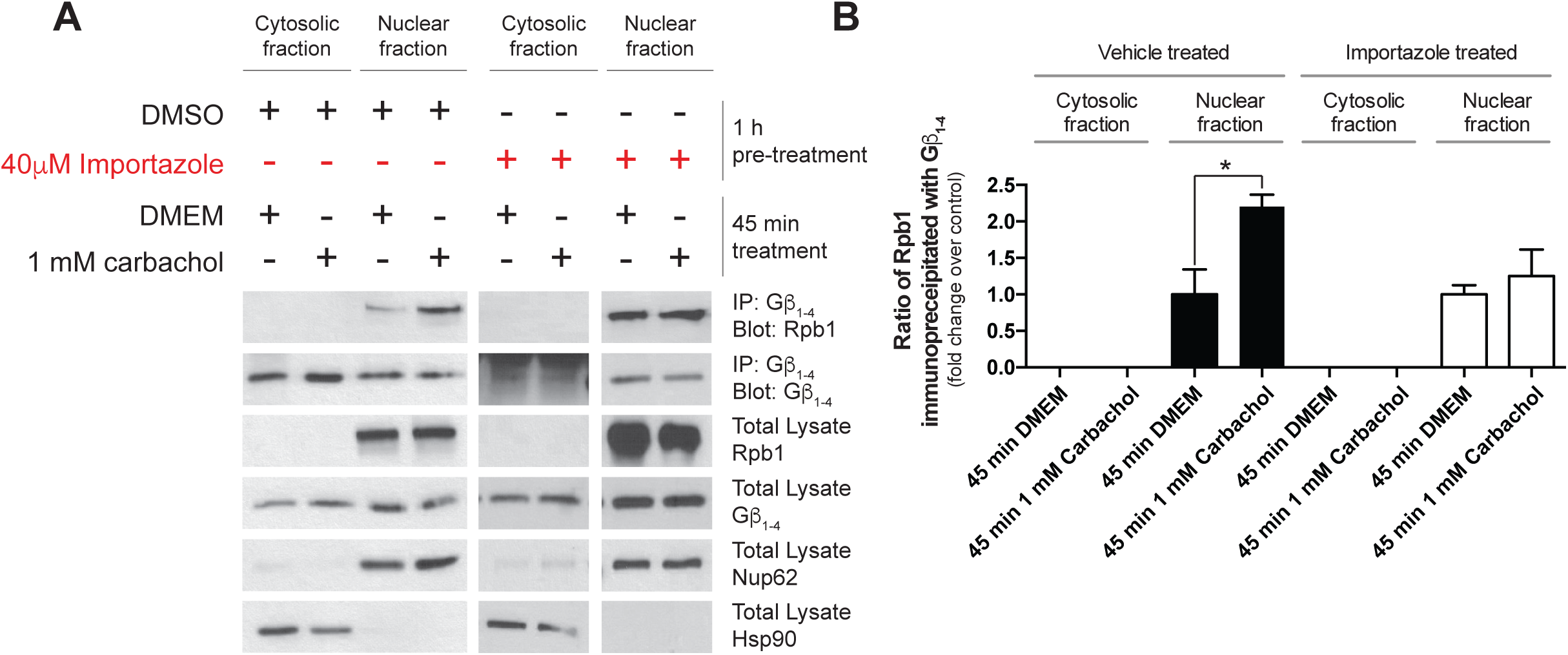
Requirement for Gβγ nuclear transport for RNAPII interaction in HEK 293 cells. **(A)** Representative experiment assessing the requirement of importin-β inhibition on the Gβγ-RNAPII interaction by sub-cellular fractionation and co-immunoprecipitation. **(B)** Densitometry-based quantification of the carbachol induced interaction and the effect of nuclear import inhibition on interaction induction. Data represents mean ± S.E.M. of three independent experiments for black bars, and two independent experiments for white bars (nuclear import inhibition conditions).

**Supplemental Figure 3.**
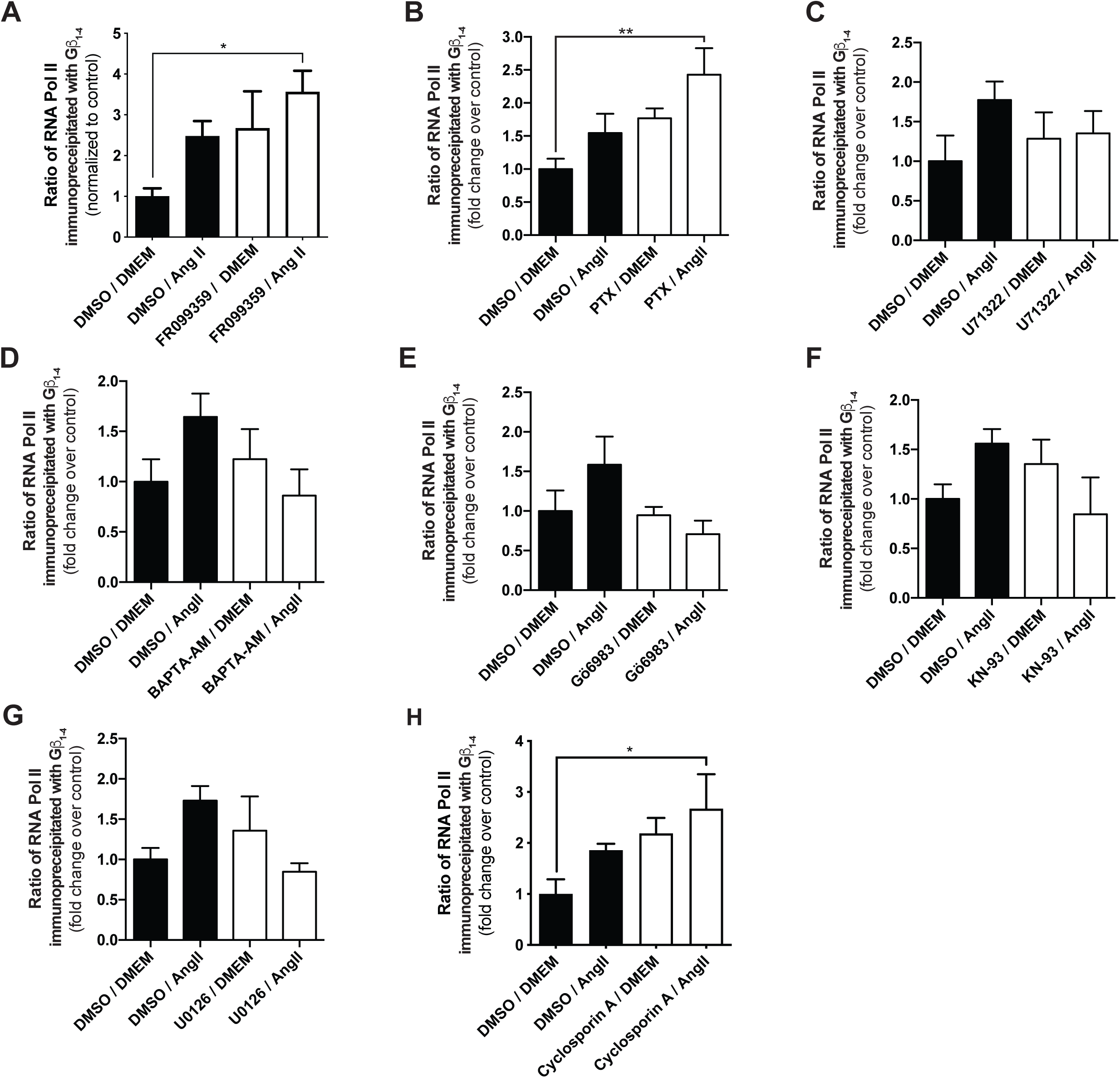
Quantitative analysis of the effects of inhibition of signalling molecules downstream of AT1R activation. **(A-H)** The relative quantities of Rpb1 co-immunoprecipitated with Gβ_1-4_ under different conditions depicted in **Figure 2** were quantified using ImageJ and normalized to DMSO/DMEM control conditions. Data shown is representative of (A) 3, (B) 6, (C) 3, (D) 4, (E) 5, (F) 5, (G) 4 or (H) 3 independent co-immunoprecipitation and western blot experiments. Data is represented as fold change over respective controls and error bars represent S.E.M. * indicates p<0.05, ** indicates p<0.01.

**Supplemental Figure 4.**
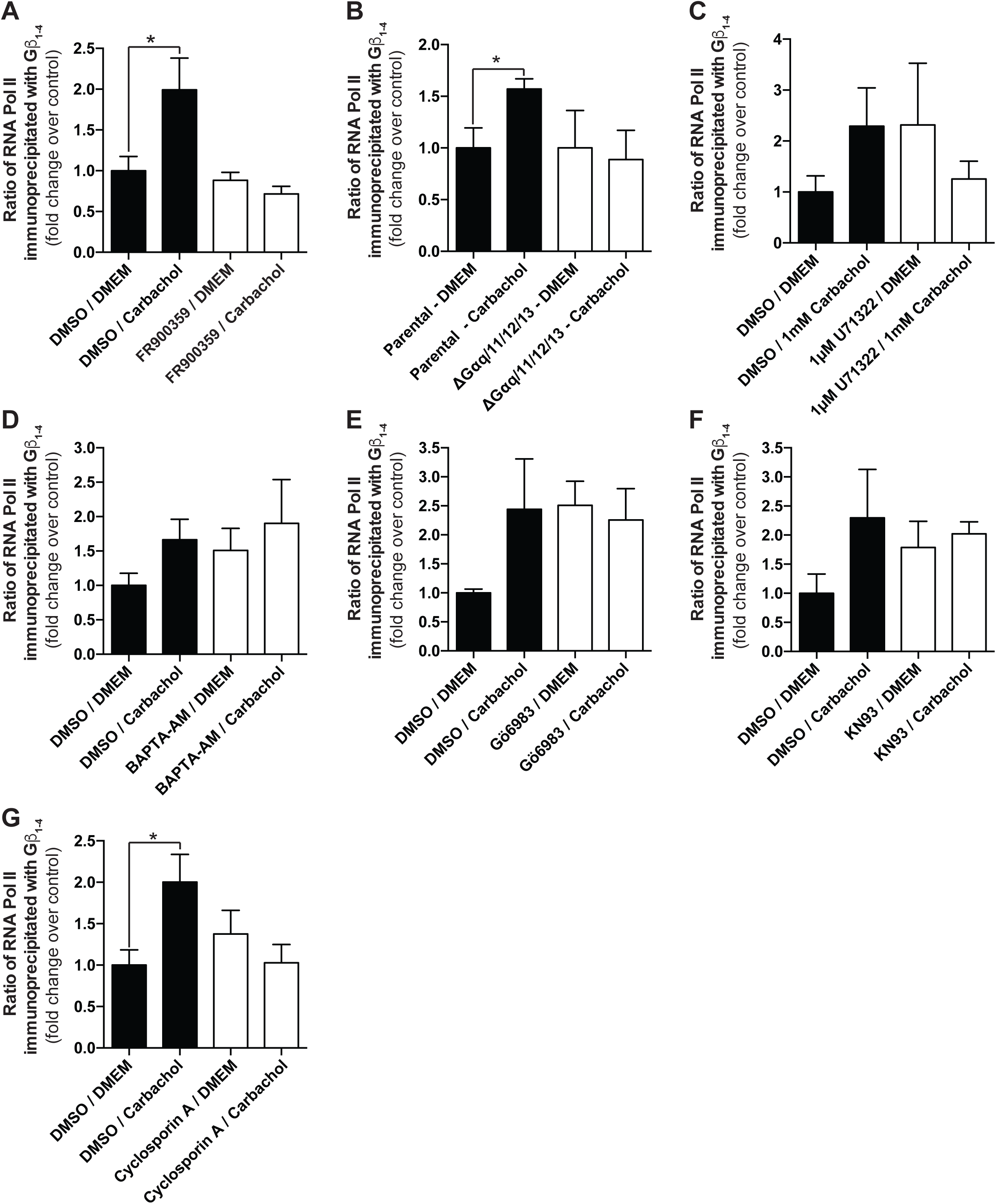
Quantitative analysis of the effects of inhibition of signalling molecules downstream of M3-mAChR activation in HEK 293 cells. **(A-G)** The relative quantities of Rpb1 co-immunoprecipitated with Gβ_1-4_ under different conditions depicted in **Figure 3** were quantified using ImageJ and were normalized to amounts pulled down in DMSO/DMEM control conditions. Data shown is representative of (A) 3, (B) 4, (C) 5, (D) 3, (E) 3, (F) 6 or (G) 4 independent co-immunoprecipitation and western blot experiments. Data is represented as fold change over respective controls and error bars represent S.E.M. * indicates p<0.05.

**Supplemental Figure 5.**
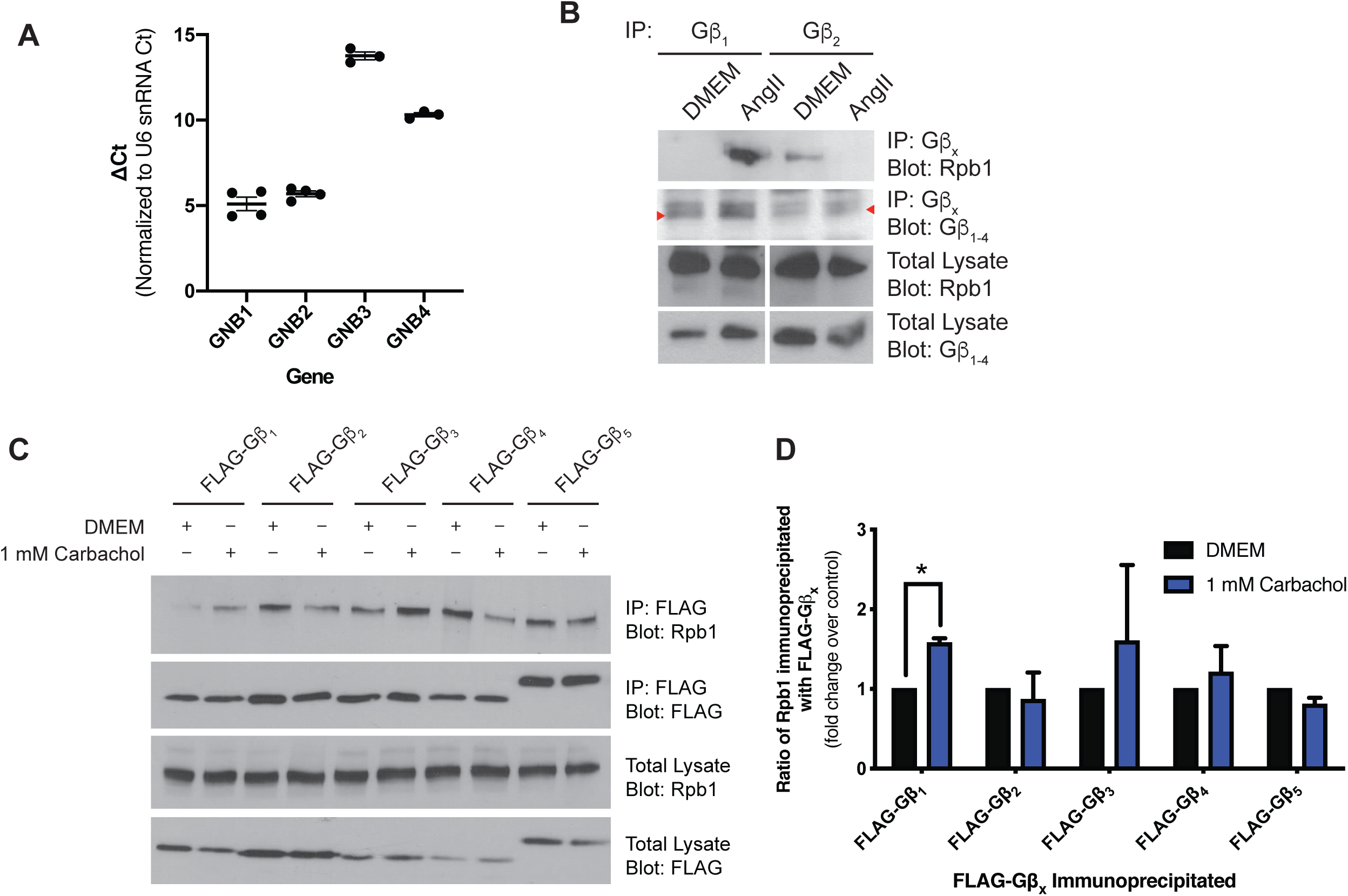
Assessment of specific Gβ subunits interacting with RNAPII upon agonist stimulation in rat cardiac fibroblasts or in HEK 293 cells. **(A)** Transcript levels for GNB1, GNB2, GNB3, and GNB4 were assessed in cardiac fibroblasts by RT-qPCR. The Ct values for each gene transcript were normalized to the house keeping U6 snRNA gene transcript for comparison. Data represents mean ± S.E.M for 3-4 independent experiments. **(B)** Gβ_1_ and Gβ_2_ were immunoprecipitated with isoform specific antibodies from cardiac fibroblasts lysates treated with 1 µM Ang II for 75 min. The amount of Rpb1 pulled down with either Gβ isoform was assessed by western blot. **(C)** Assessment of specific FLAG-tagged Gβ isoforms interaction with Rpb1 under conditions of M3-mAChR stimulation with carbachol in HEK 293 cells. The amount of Rpb1 interacting with each Gβ isoform was assessed by western blot following FLAG immunoprecipitation. **(D)** Densitometry-based quantification of the ratio of Rpb1 co-immunoprecipitated with the indicated FLAG-tagged Gβ subunit. The ratio of Rpb1 to FLAG-Gβ_x_ immunoprecipitated was determined and normalized to fold change over DMEM treatment. Data represents mean ± S.E.M for four independent replicates; * indicates p<0.01

**Supplemental Figure 6.**
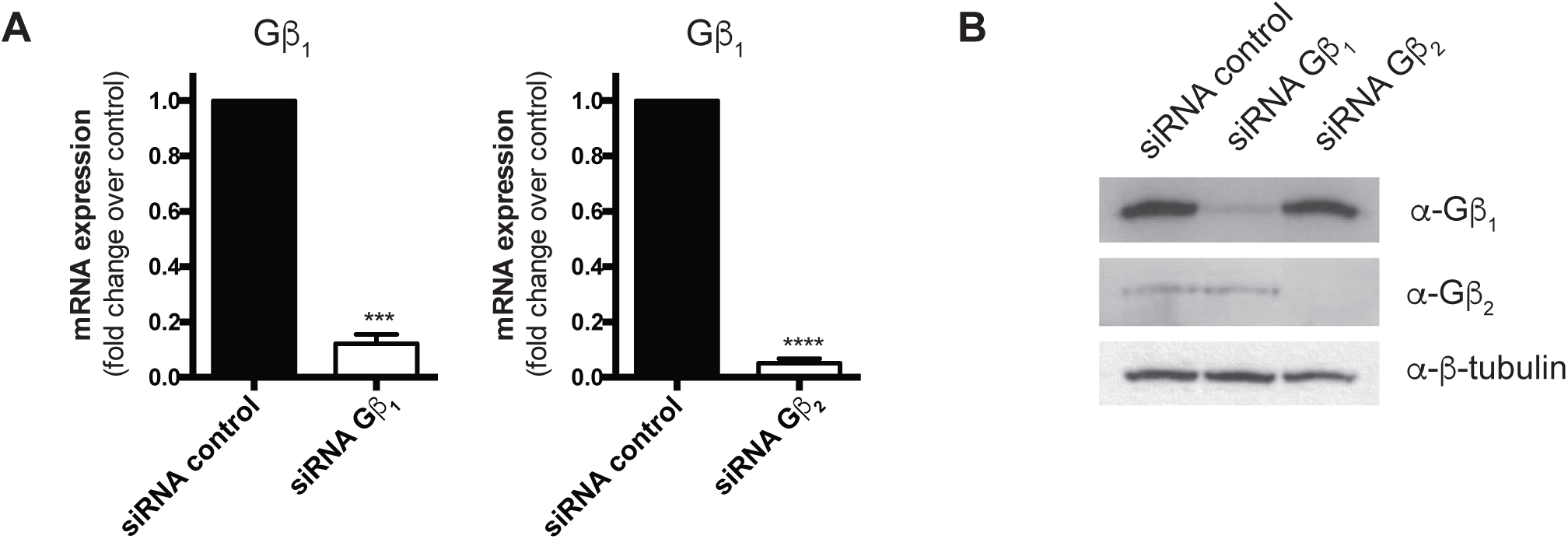
Validation of RNAi knockdown of Gβ_1_ and Gβ_2_. Validation of Gβ_1_ and Gβ_2_ mRNA **(A)** and protein **(B)** knockdown with siRNA in rat neonatal cardiac fibroblasts. Rat neonatal cardiac fibroblasts were transfected with 50 nM siRNA control, Gβ_1_ or Gβ_2_ for 72 hours, serum-deprived for 12 h and RNA or protein collected as described in *Methods*. Data in (A) represents mean ± S.E.M for four independent experiments; * Ct values were normalized to the housekeeping U6 snRNA transcript and fold change over siRNA control determined using the 2^-ΔΔCt^. ** indicates p<0.001 and **** indicates p<0.0001.

**Supplemental Figure 7.**
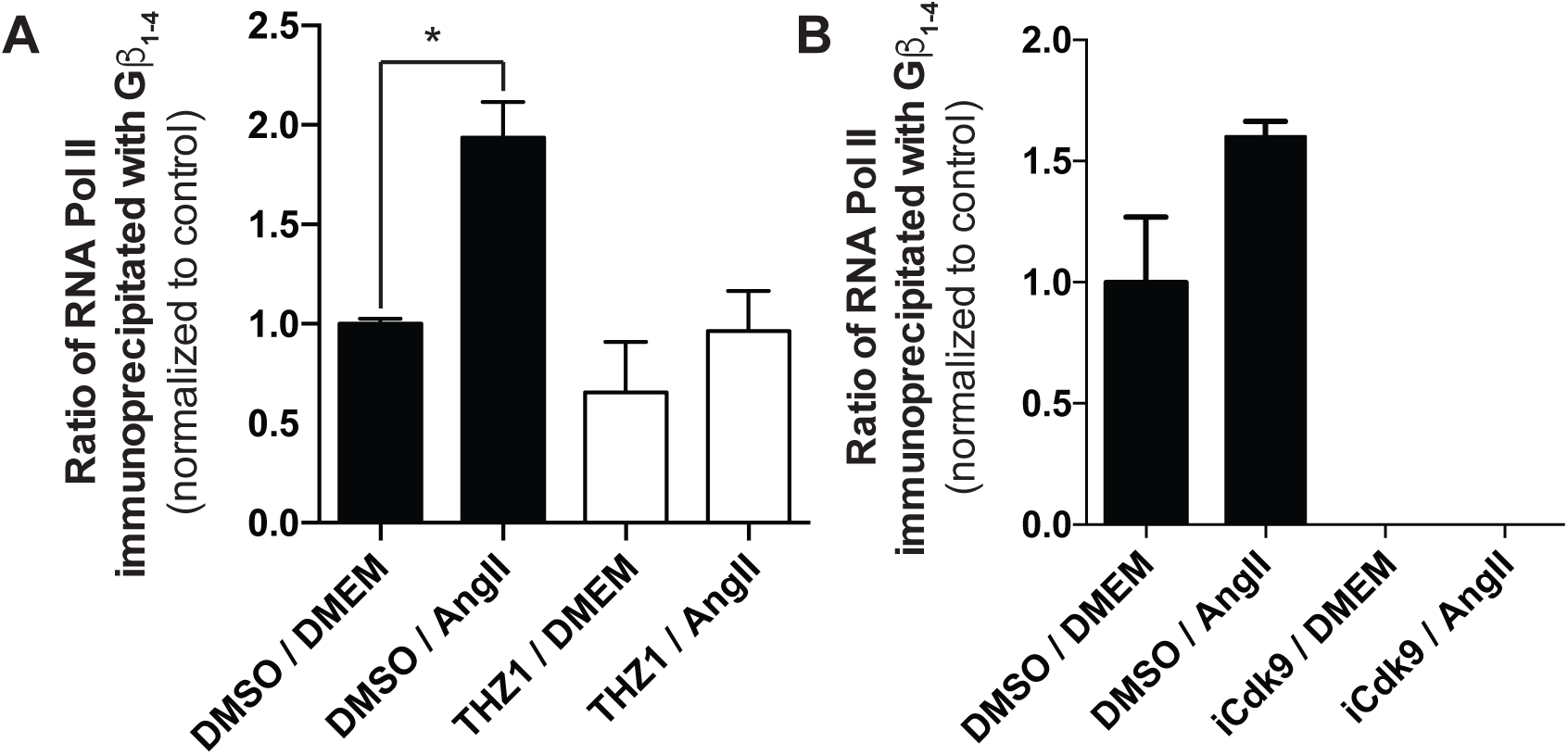
Quantitative analysis of the effect of transcriptional regulator inhibition on the Gβγ-RNAPII interaction in cardiac fibroblasts. **(A-B)** The relative quantities of Rpb1 co-immunoprecipitated with Gβ_1-4_ under conditions depicted in Figure 5A (THZ1) and B (iCdk9) were quantified and normalized to DMSO/DMEM control conditions. Data shown is representative of between three to six independent co-immunoprecipitation and western blot experiments. Data represents mean ± S.E.M. * indicates p<0.05.

**Supplemental Figure 8.**
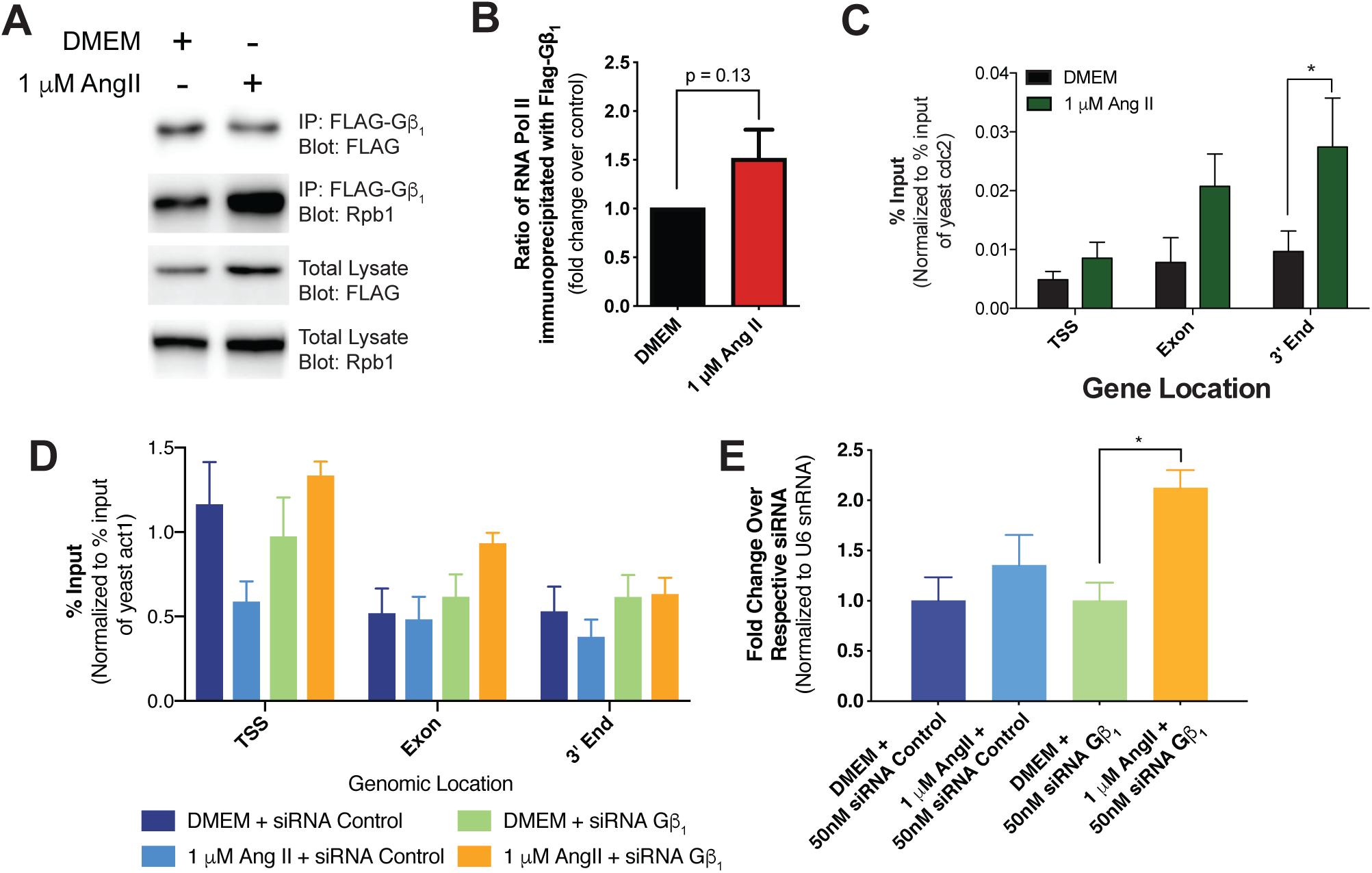
Validation of heterologously expressed FLAG-tagged Gβ_1_ in rat neonatal cardiac fibroblasts. **(A)** Assessment of Rpb1 co-immunoprecipitated with FLAG-Gβ_1_ following 75 min treatment of 1 µM Ang II in rat neonatal cardiac fibroblasts. Cardiac fibroblasts were transduced with AAV1-FLAG-Gβ_1_ prior to treatment with 1 µM Ang II. **(B)** Densitometry-based quantification of the ratio of Rpb1 co-immunoprecipitated with FLAG-Gβ_1_. The ratio of Rpb1 to FLAG-Gβ_1_ was calculated and normalized as fold change over DMEM condition. Data is represented as mean ± S.E.M for four independent experiments. Assessing changes in FLAG-Gβ_1_ **(C)** or **(D)** Rpb1 occupancy along Ctgf following 75 min treatment with 1 µM Ang II. FLAG-Gβ_1_ or Rpb1 was immunoprecipitated from crosslinked and sonicated chromatin, DNA purified and quantified by qPCR. Data is represented as mean ± S.E.M for 4-6 independent experiments, * indicates p<0.05. **(E)** Validation of Ctgf gene expression with primers distinct from those used in the Qiagen RT^2^ Profiler^TM^ PCR array. Data represents mean ± S.E.M for four independent experiments, * indicates p<0.05.

**Supplemental Figure 9.**
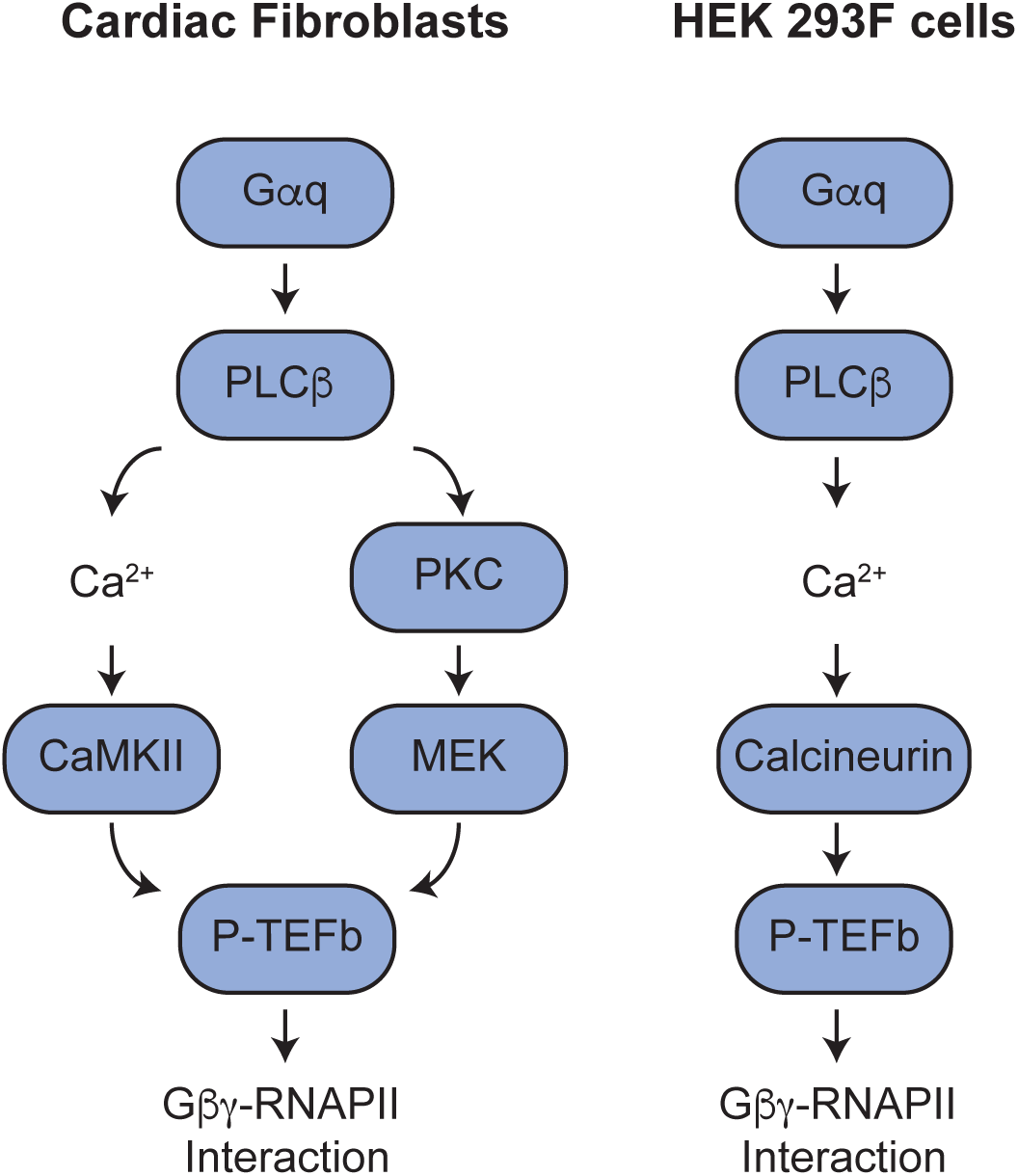
Schema summarizing signalling events regulating the agonist induced Gβγ interaction with RNAPII. Signalling cascade downstream of AT1R in cardiac fibroblasts or M3 muscarinic receptors in HEK 293F cells regulating the interaction. Signalling pathways were determined by assessing Gβγ-RNAPII interactions by co-immunoprecipitation and western blot as shown in **Figure 2** and Supplemental Figure 3 for cardiac fibroblasts and **Figure 3** and Supplemental Figure 4 for HEK 293F cells.

**Supplemental Table 1.** Summary of fibrosis RT-qPCR array results. This table summarizes gene expression changes measured using the Qiagen RT^2^ Profiler^TM^ PCR Array at 75 min and 24 h Ang II stimulation. Genes were considered to have altered expression with fold changes ≥ 1.5 or ≤ 0.5 compared to DMEM/siRNA control conditions at the respective time point. In parenthesis are the number of genes with a significant (p<0.05) change in expression compared to DMEM/siRNA control at the respective time point. Two-way ANOVA followed by post-hoc t-test comparisons with Bonferroni correction was performed for each gene individually. Data is representative of three independent biological replicates.

| Time | siRNA | Treatment | Upregulated | Downregulated |
| --- | --- | --- | --- | --- |
| 75 min | Control | DMEM | 0 | 0 |
| | | 1 μM Ang II | 7(5) | 0 |
|  | Gnb1 | DMEM | 4(1) | 1 |
| | | 1 μM Ang II | 10(6) | 2 |
| 24 h | Control | DMEM | 0 | 0 |
| | | 1 μM Ang II | 37(7) | 0 |
|  | Gnb1 | DMEM | 26(2) | 1 |
| | | 1 μM Ang II | 53(13) | 0 |
|  | Gnb2 | DMEM | 7 | 1 |
| | | 1 μM Ang II | 44(11) | 0 |

**Supplemental Table 2.**
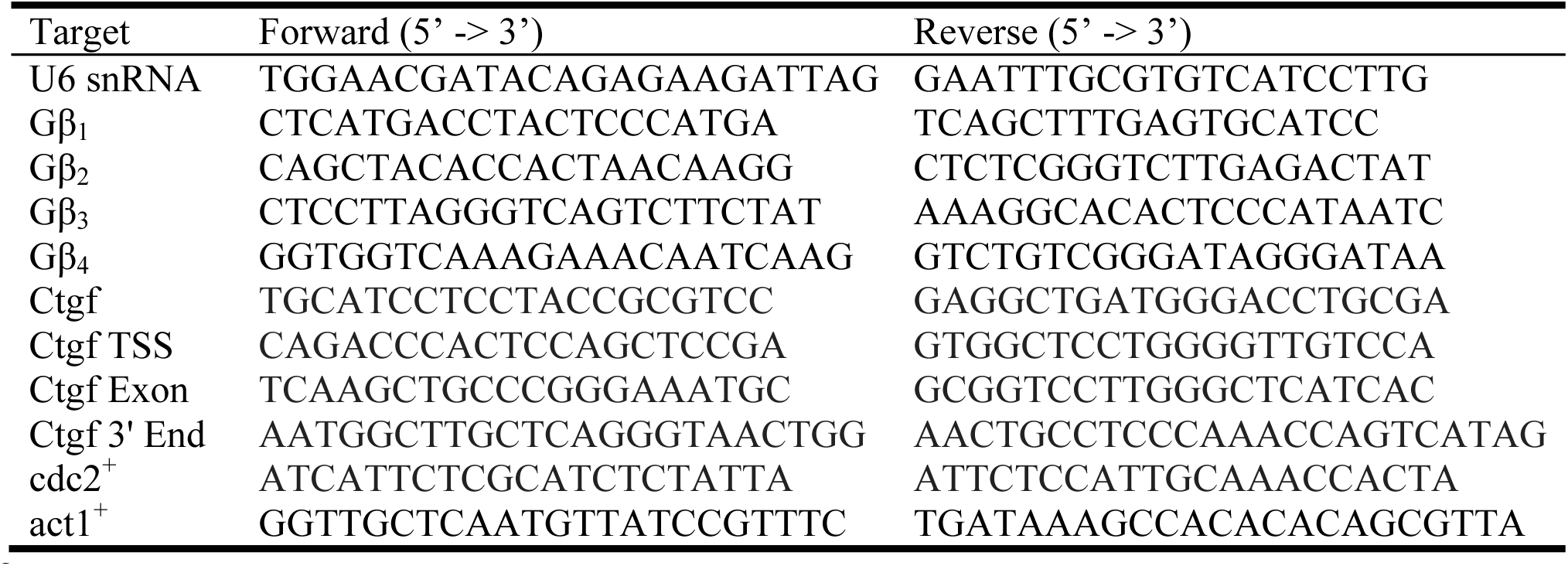
List of primers used to assess gene expression by RT-qPCR and ChIP-qPCR in cardiac fibroblasts. Forward and reverse primers were used at a concentration of 300 nM for each qPCR reaction. Primer sequences were designed using NCBI’s Primer-BLAST tool and validated by analysis of standard curve qPCR assays performed in-house.

